# Structural architecture of TolQ-TolR inner membrane protein complex from opportunistic pathogen *Acinetobacter baumannii*

**DOI:** 10.1101/2024.06.19.599759

**Authors:** Elina Karimullina, Yirui Guo, Hanif M. Khan, Tabitha Emde, Bradley Quade, Rosa Di Leo, Zbyszek Otwinowski, D. Tieleman Peter, Dominika Borek, Alexei Savchenko

## Abstract

Gram-negative bacteria harness the proton motive force (PMF) within their inner membrane (IM) to uphold the integrity of their cell envelope, an indispensable aspect for both division and survival. The IM TolQ-TolR complex is the essential part of the Tol-Pal system, serving as a conduit for PMF energy transfer to the outer membrane. Here we present cryo-EM reconstructions of *Acinetobacter baumannii* TolQ in apo and TolR- bound forms at atomic resolution. The apo TolQ configuration manifests as a symmetric pentameric pore, featuring a trans-membrane funnel leading towards a cytoplasmic chamber. In contrast, the TolQ-TolR complex assumes a proton non-permeable stance, characterized by the TolQ pentamer’s flexure to accommodate the TolR dimer, where two protomers undergo a translation-based relationship. Our structure-guided analysis and simulations support the rotor-stator mechanism of action, wherein the rotation of the TolQ pentamer harmonizes with the TolR protomers’ interplay. These findings broaden our mechanistic comprehension of molecular stator units empowering critical functions within the Gram-negative bacterial cell envelope.

**Teaser:** Apo TolQ and TolQ-TolR structures depict structural rearrangements required for cell envelope organization in bacterial cell division.

## Introduction

The cell envelope is a complex barrier system that shields bacterial cytoplasm and facilitates bacterial survival in a rapidly-changing environment. In Gram-negative bacteria, the canonical structure of the cell envelope consists of a peptidoglycan layer (PG) sandwiched between outer and inner bilayer lipid-based membranes. The outer membrane (OM) composition is asymmetric with the inner phospholipid layer and the outer layer made of lipopolysaccharide. Lacking systems to power active processes in OM, Gram-negative bacteria have evolved protein complexes spanning the cell envelope and connecting OM with the inner membrane (IM). Such systems allow for the energy generated by the proton motive force (PMF) in the IM to be used to drive reactions at OM. One such system is called Tol-Pal, in which the energy-transducing IM complex composed of TolQ, TolR, and TolA proteins, using PMF energy to modulate interactions of TolA with components of the TolB-Pal OM complex (*1–5*) (Fig. 1A). The Tol-Pal system is involved in the remodeling of peptidoglycan at cell division sites and is implicated in other cell division steps involving OM invagination (*6–8*). Indispensable for Gram-negative cell envelope Tol-Pal protein complex is targeted by bacterial toxins - colicins and is a frequent entry point for filamentous phages (*9, 10*), which exploit pulling- in mechanical force generation by TolQ-TolR-TolA complex.

**Fig. 1.**
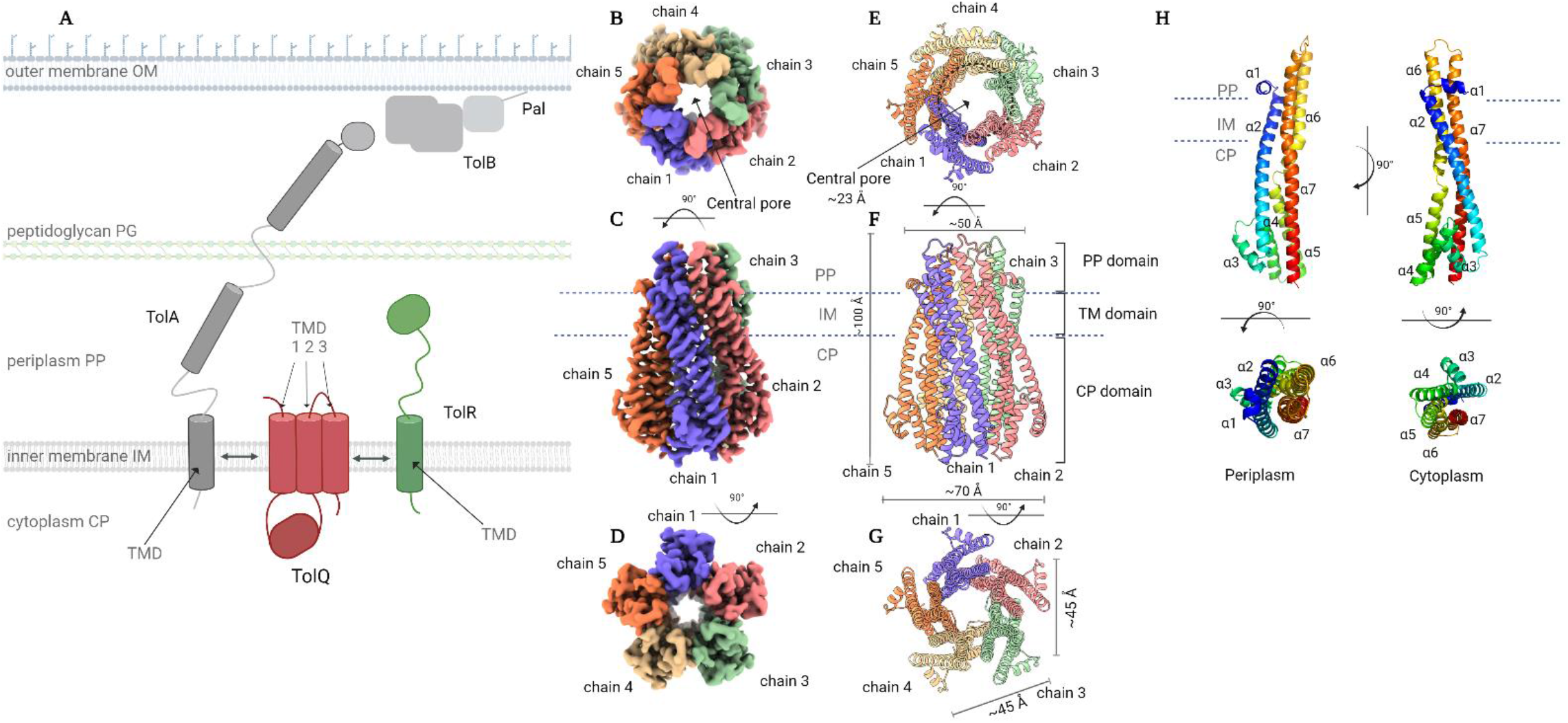
Architecture and Topology of the TolQ Cryo-EM structure. **(A)** Schematic of the Tol-Pal system for energy transduction. **(B)** Periplasmic side view of the cryo-EM map of the TolQ pentamer. **(C)** Side view of the cryo-EM map of the TolQ. **(D)** Cytoplasmic view of the cryo-EM map of the TolQ pentamer. **(E)** Periplasmic side view of the cartoon model of the TolQ pentamer. **(F)** Side view of the cartoon model of the TolQ pentamer. **(G)** Cytoplasmic side view of the cartoon model of the TolQ pentamer. **(H)** Cartoon representation of the TolQ monomer.

Previous studies of Tol-Pal complex components have established some characteristics of individual components of this system. For TolA protein, three domains with distinct functions were identified. The N-terminal domain of TolA is embedded into IM and is followed by the long linker domain and C-terminal domains; both the linker domain and the C-terminal domains are shown to be responsible for interactions with various components of OM or proteins associated with OM (*1*), including TolB (*11*). TolR is a small protein that is predicted to contain a single N-terminal α-helical transmembrane domain (TMD) followed by a periplasmic domain that is expected to undergo a structural transition to make contacts with the peptidoglycan (PG) layer (*12, 13*). The TolQ has been described as a multipass transmembrane protein containing three TMDs and a large cytoplasmic insertion domain located between the first and second TMDs (*13–16*) (Fig. 1A). Mutations in IM segments of all three proteins result in severe phenotypical aberrations, which emphasizes the importance of interactions in TolQ-TolR-TolA complex (*1, 3, 17–21*). In *Escherichia coli* the point mutations in TolQ transmembrane segments alter the cell envelope integrity (*14*) and result in RNase I leakage and formation of outer membrane vesicles (*1*). Similarly, point mutations in TolQ affected the permeability of the cell envelope in *Salmonella typhimurium* (*22*) and bacteremia-causing *Erwinia chrysanthemi* (*23, 24*), sensitizing these bacteria to bile salts and decreasing their survival. While ΔtolQ is not lethal in *E. coli*, it causes reduced O7 lipopolysaccharide expression, lower motility, and deficiency in colonization (*25, 26*). ΔtolQ mutants of *Edwardsiella ictalurid* (*27*), *S. typhimurium* (*22*) and *S. holeraesuis* (*28*) also demonstrated decreased virulence.

The established role of the Tol-Pal system in the virulence and survival of Gram-negative bacteria during pathogenesis makes it an attractive target for antibacterial intervention and is driving intensive mechanistic studies of this system. These studies include the characterization of the periplasmic domain of TolA protein from several bacterial species including *Pseudomonas aeruginosa* (*29*), *E. coli* (*30*), and *Vibrio cholerae* (*31*).

Additionally, the structure of the corresponding domain of *E. coli* TolA has been determined in complex with the N-terminal domain of g3p phage (*30*) and with the colicin A (*32*). However, the structure of full-length TolA remains uncharacterized. Similarly, the structures of the putative periplasmic TolR fragments have been determined for *Haemophilus influenzae* (*33*) and *E. coli* (*12*) with NMR and X-ray crystallography, respectively. However, the membrane-bound TolR segment has not been experimentally characterized at atomic resolution. TolQ is the only component of the TolQ-TolR-TolA subcomplex for which no experimentally derived atomic structures have been available.

Recently, a cryo-EM reconstruction of the TolQ-TolR sub-complex from *E. coli* was reported (*34*). It revealed 5:2 stoichiometry for the TolQ-TolR subcomplex with transmembrane helices of TolR dimers located inside the TolQ pentamer. However, the resolution of obtained data (the 4.3 Å overall resolution of reconstruction for the complex and 4.7 Å resolution for the TolR component) did not allow for modeling TolQ-TolR interactions.

The model for the molecular mechanism of the Tol-Pal system has been proposed based on the flagellar motor system MotA-MotB and nutrient transport system ExbB-ExbD, the components of which share sequence similarly with TolQ and TolR (*1, 11, 35*). Recent cryo-EM and X-ray crystallographic studies identified 5:2 stoichiometry for both systems, with five monomers of MotA/ExbB surrounding the transmembrane part of two MotB/ExbD molecules (*36–39*). Based on sequence similarities with MotA-MotB and ExbB-ExbD systems, the TolQ-TolR is expected to operate as the stator-rotor motor (*40*). In such a system, stators are stationary parts of the rotary motors, while energy provided to the motor generates torque applied to the rotor. This model has been best demonstrated in the MotA-MotB system, where the MotB acts as a stator. However, it is less certain whether the homologous components of the other two systems adopt the same mode of action.

Here, we determined a 3.02 Å cryo-EM SPR structure of the full-length TolQ in apo form and a 3.34 Å cryo-EM structure of the TolQ-TolR complex from *Acinetobacter baumannii*. *A. baumannii* is one of the most dreaded and clinically important Gram-negative ESKAPE (*Enterococcus faecium*, *Staphylococcus aureus*, *Klebsiella pneumoniae*, *Acinetobacter baumannii*, *Pseudomonas aeruginosa*, and *Enterobacter* spp.) pathogens causing a broad-spectrum nosocomial infection mainly among patients admitted to intensive care units (*41–44*).

Our findings offer experimentally derived, high-resolution structural models of both TolQ and the TolQ-TolR complex. These models provide insights into potential mechanisms of proton translocation and force generation within the Tol-Pal system, serving as a reference for analyzing force generation in other 5:2 stator-rotor systems. Additionally, they provide the molecular framework for the rational design of inhibitors against a vital molecular complex conserved across Gram-negative bacteria.

## Results

### Molecular architecture of TolQ pentamer

As an initial step in the structural characterization of Tol-Pal system components in *Acinetobacter baumannii*, we pursued the structural characterization of the TolQ protein. The corresponding gene was cloned and recombinantly expressed in *E. coli* (see Material and Methods for details). TolQ protein was purified from the membrane fraction of the cell lysate by the combination of affinity and size exclusion chromatography (fig. S1A-D). Once purified, TolQ protein was reconstructed using amphipol polymers. According to mass photometry analysis (fig. S1D), solubilized TolQ protein formed homogeneous particles having mass higher than 100 kDa thus amenable to analysis by cryo-EM SPR.

The TolQ sample was biochemically and conformationally homogenous and generated well-behaving cryo-EM grids. Upon data acquisition, 2D classification of selected particles revealed clear features of pentameric stoichiometry (fig. S2A). After 3D reconstruction with C5 symmetry, we obtained a 3.02 Å molecular density map (fig. S2, B-D; table S2). The analysis of the map revealed five molecules of TolQ forming a pentameric assembly of ∼100 Å long with ∼70 Å diameter (Fig. 1, B-D), stabilized by a ring of A8-35 amphipol visible at lower contour density map at TMD region (fig. S3, A-C). To facilitate the comparative analysis, we named the chains in TolQ pentamer following nomenclature reported in previous studies for the TolQ homolog MotA (*38*) with chain 1 (See Fig. 1, B-F) corresponding to chain A in PDB structure 6YKM and so on.

According to our analysis, TolQ protomers contain seven α-helices (Fig. 1H). The N-terminal α1 helix is positioned along the periplasmic surface of IM. The α2 helix spans the IM forming the first transmembrane helical region (TMH1) and extends into the cytoplasm. α2 helix is followed by two cytoplasmic, relatively short, α3 and α4 helices.

Cytoplasmic α5 helix loops back to the IM and is followed by α6 helix, corresponding to TMH2, which spans the membrane back to periplasmic space. The C-terminal and the longest α7 helix spanning ∼100 Å, forms TMH3. All helices are connected via relatively short loops. In our structure, the outer surfaces of TMH1 and TMH2 interact directly with amphipol while TMH3 forms the inner surface of the TolQ central pore. This central tunnel has a circular opening of ∼23 Å in diameter at the periplasmic side (Fig. 1E). The surface buried by interactions between two TolQ monomers is 1229-1234 Å^2^ per each interface (table S1), with both periplasmic and cytoplasmic segments of TolQ also contributing to interactions (Fig. 1, E-G). A small gap exists between adjacent protomers on the periplasmic side (fig. S4).

According to Foldseek server (*45*), our *Ac*TolQ protomer structure is the most similar to the structure of ExbB from *E. coli* (PDB: 6TYI) and *Serratia marcescens* (PDB:7AJQ) with RMSD of 1.60 Å over 193 Cα and 2.39 Å over 159 Cα atoms, respectively. The second closest structure is flagellar stator protein MotA from *Aquifex aeolicus* and *Bacillus subtillis*, although this protein’s protomers contain an additional cytoplasmic α- helix lacking in TolQ (*38, 46*). Both MotA (PDB:8GQY) and ExbB (PDB:6TYI and/or 6YE4) form pentameric symmetrical structures with the pore entrance from periplasmic side, similar to TolQ apo structure presented here (fig. S5, A-C).

### Cryo-EM structure of TolQ-TolR complex

For structural characterization of *A. baumannii* TolQ protein in complex with TolR, TolQ and TolR genes were cloned and co-expressed using a tandem promoter vector system in *E. coli* (see Material and Methods for details). After purification using affinity and size exclusion chromatography, the fraction corresponding to the TolQ-TolR complex (fig. S1, B and C) was further analyzed by mass photometry. This analysis suggested that TolQ-TolR complex was larger than TolQ pentamer by ∼30 kDa, which would correspond to the TolR dimer (fig. 1D). Cryo-EM SPR of TolQ-TolR complex confirmed 5 to 2 stoichiometry (Fig. 2, A-H). We will refer to this complex hereafter as *Ac*TolQ-TolR. The reconstruction of *Ac*TolQ-TolR reached overall resolution of 3.34 Å with C1 symmetry imposed (fig. S5, A-F and fig. S2).

**Fig. 2.**
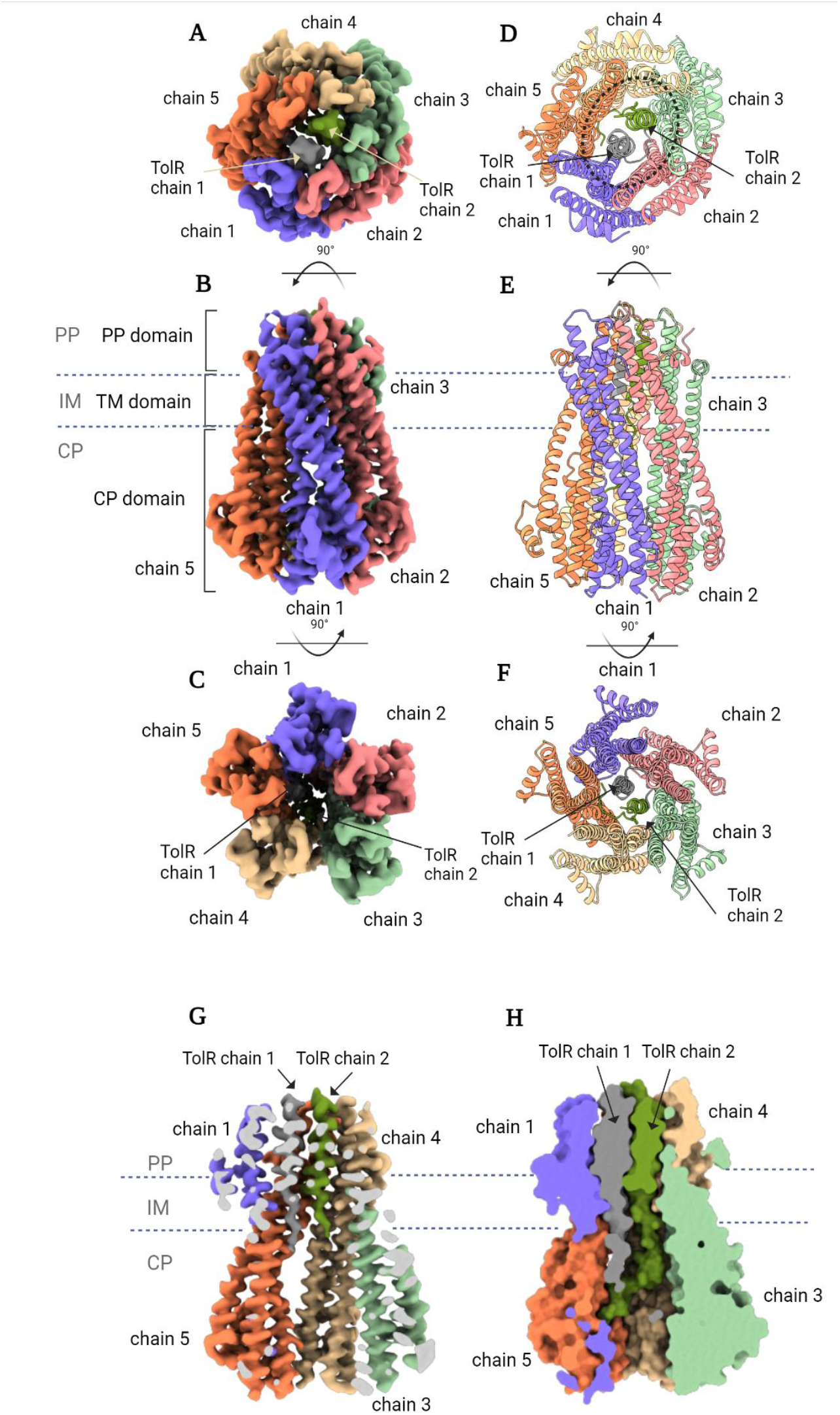
Architecture and Topology of the TolQ-TolR complex Cryo-EM structure. **(A)** Periplasmic side view of the cryo-EM map of the TolQ-TolR 5:2 complex. **(B)** Side view of the cryo-EM map of the TolQ-TolR 5:2 complex. **(C)** Cytoplasmic view of the cryo-EM map of the cryo-EM map of the TolQ-TolR 5:2 complex. **(D)** Periplasmic side view of the cartoon model of the TolQ-TolR 5:2 complex. **(E)** Side view of the cartoon model of the TolQ-TolR 5:2 complex. **(F)** Cytoplasmic side view of the cartoon model of the TolQ-TolR 5:2 complex. **(G)** Side view of Cryo-EM map of TolQ-TolR complex with the front of the complex removed. **(H)** Side view of TolQ-TolR complex surface representation of the model with the front of the complex removed.

Comparison of TolQ protomers conformations in apo and TolR-bound structures using PyMol revealed modest rearrangements reflected in very low RMSD value (0.50 to 0.82 Å) between TolQ protomers in both structures over 215 Cα atoms (Fig. 3A). However, superimposing tertiary structures, we observed a shift in the overall position of Chain 1 in AcTolQ versus AcTolQ-TolR structures to maximum of 9.32 Å at periplasmic side of the complex (Fig. 3, B and C). As a result, TolQ pentamer’s central pore in the complex structure underwent the shape change from round to oval (Fig. 2D) with ∼30 Å in the widest diameter accommodating the TMHs of TolR dimer inside the pore (Fig. 2, G and H and Fig. 3, B and C). The average variation for superimposed TolQ pentamers in apo and TolR-bound structures was 1.91 Å. The analysis of interface areas between protomers calculated by PISA highlights the asymmetry between apo and bond TolQ forms in more detail (table S1). While interface area levels are equal (∼1200 Å^2^) between TolQ protomers in the apo form, the interface area between TolQ chains varies greatly (755- 1439 Å^2^) in the AcTolQ-TolR complex with C1 symmetry. The largest interface areas are between chains 1-2, 2-3, and 4-5 of TolQ. The wider gap between chains 3-4 and 5-1 results in smaller interface areas 755 and 823 Å^2^, respectively (table S1).

**Fig. 3.**
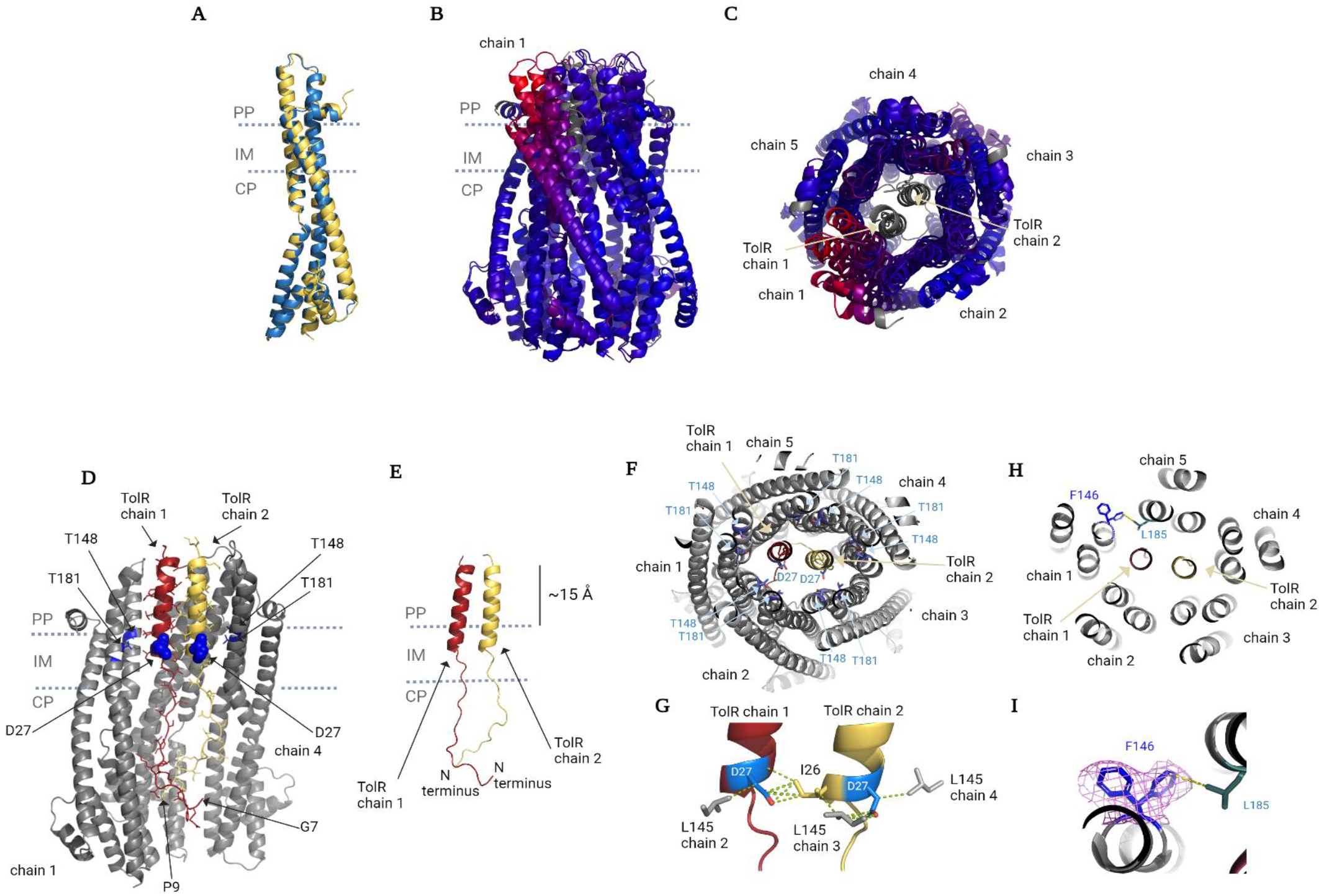
Arrangements of TolQ and TolR protomers in the complex. **(A)** The superimposed TolQ protomers from TolQ (blue) and TolQ-TolR (yellow) Cryo-EM structures. **(B)** The superimposed pentameric TolQ and 5:2 TolQ-structures colored by RMSD, side view. Dark blue is good alignment, higher deviations are red. Residues not used for alignment are colored white **(C)** The superimposed pentameric TolQ and 5:2 TolQ-TolR structures colored by RMSD, periplasmic side view. Dark blue is good alignment, higher deviations are red. Residues not used for alignment are colored white. **(D)** Cartoon representation of TolR TMD and N-terminus solved structure relative to the inner membrane, side view. **(E)** Co- localization of functionally important TolR residues (blue spheres) and its surrounding environment made by TolQ residues (blue cartoon and sticks representation), side view. TolQ chain 2 is eliminated for clarity **(F)** Co- localization of functionally important TolR residues, and their surrounding environment made by TolQ, periplasmic view with the periplasmic part cut off for clarity. **(G)** Key TolR residues D27 (blue) interactions within 3.0 - 3.8 Å distances. **(H)** TolQ chain 1 F146 side chain double conformation in TolQ-TolR structure, slab view from the periplasm. **(I)** Zoomed TolQ chain 1 F146 side chain double conformation in TolQ-TolR structure with electron density depicted in magenta.

*Ac*TolQ-TolR structure revealed the position of the N-terminal portion of TolR, which, according to our knowledge, has evaded to be experimentally visualized before. The N-terminal segments of both TolR protomers localize to the cytoplasmic chamber formed by TolQ pentamer and do not have a defined secondary structure for the region prior to the transmembrane helix (TMH) that starts at the residue Tyr^25^. TolR N-terminal segments corresponding to residues Gly^7^-Ser^45^ and Phe^9^-Ser^45^ at the N-terminus of both TolR protomers are well resolved in obtained complex structure. Such positioning of TolR N-termini inside the TolQ pentamer in our model contradicts the report that proposed for this portion of TolR to be exposed to cytosol (*47*). This orientation positions the resolved N-terminus of TolR which is composed by mostly positively charged residues against the large electronegative chamber formed by TolQ pentamer (Fig. 3E and fig. S8C-F).

Following the segment lacking secondary structure, corresponding to the residues Tyr^25^ to Ser^45^ each TolR protomer forms the α-helix spanning the pore of TolQ pentamer and extending 15 Å beyond IM into the periplasmic side (Fig. 3D and fig. S7). The portion of TolR inside TolQ pentamer is accommodated by the oval shaped periplasmic side of the TolQ central pore which is lined with mostly negatively charged residues facing the periplasm side (fig. S8, A and B). The interface between TMHs in the TolR dimer buries an area of ∼600 Å^2^. This interface is almost exclusively hydrophobic (fig. S8, C-F) with six and eight residues from TolR chain 1 and chain 2, respectively, contributing over twenty non-bonding interactions (fig. S9). The N-terminal segment of TolR is connected to the C-terminal periplasmic domain via a flexible linker formed by residues Gly^46^ to Ala^66^, which are not visible in obtained complex structure. The C-terminal portion of TolR is referred to in the literature as the peptidoglycan binding domain (PGBD) (*12*). The PGBD’s shape was determined to a very low resolution in *Ac*TolQ-TolR indicating significant movement of the domain (fig. S6A and S7). The low-resolution map obtained for this portion of TolR agrees with previously determined crystal and NMR structures of this domain (*12, 33*), which can be confidently modeled into it (data not shown).

Overall, the general architecture of *Ac*TolQ-TolR complex is similar to the architecture of analogous sub-complexes from ExbB-ExbD and MotA-MotB systems (*37, 38*). However, the α-helical regions of two TolR protomers in the *Ac*TolQ-TolR complex are related by translational symmetry rather than by rotational symmetry observed in other systems (Fig. 3, E-G).

### Analysis of TolQ-TolR interactions

Atomic resolution cryo-EM SPR enabled detailed analysis of interactions between *A. baumanii* TolQ and TolR proteins and interpretation of these interactions in context of Tol-Pal system cellular activity based on results of previously conducted biochemical or genetic experiments (*1, 15, 17–19, 23, 47*). Specifically, the analysis of TolQ-TolR systems in *E. coli* and *S. typhimurium* identified a number of mutations critical for the function and interactions between these proteins (*1, 15, 17–19, 23, 47*). Since the primary sequences of these proteins show significant similarity with *A. baumannii* TolQ and TolR (46 % of sequence ID in case of TolQ) we investigated the role of corresponding residues in *Ac*TolQ5-TolR2 structure (Fig. 4A).

**Fig. 4.**
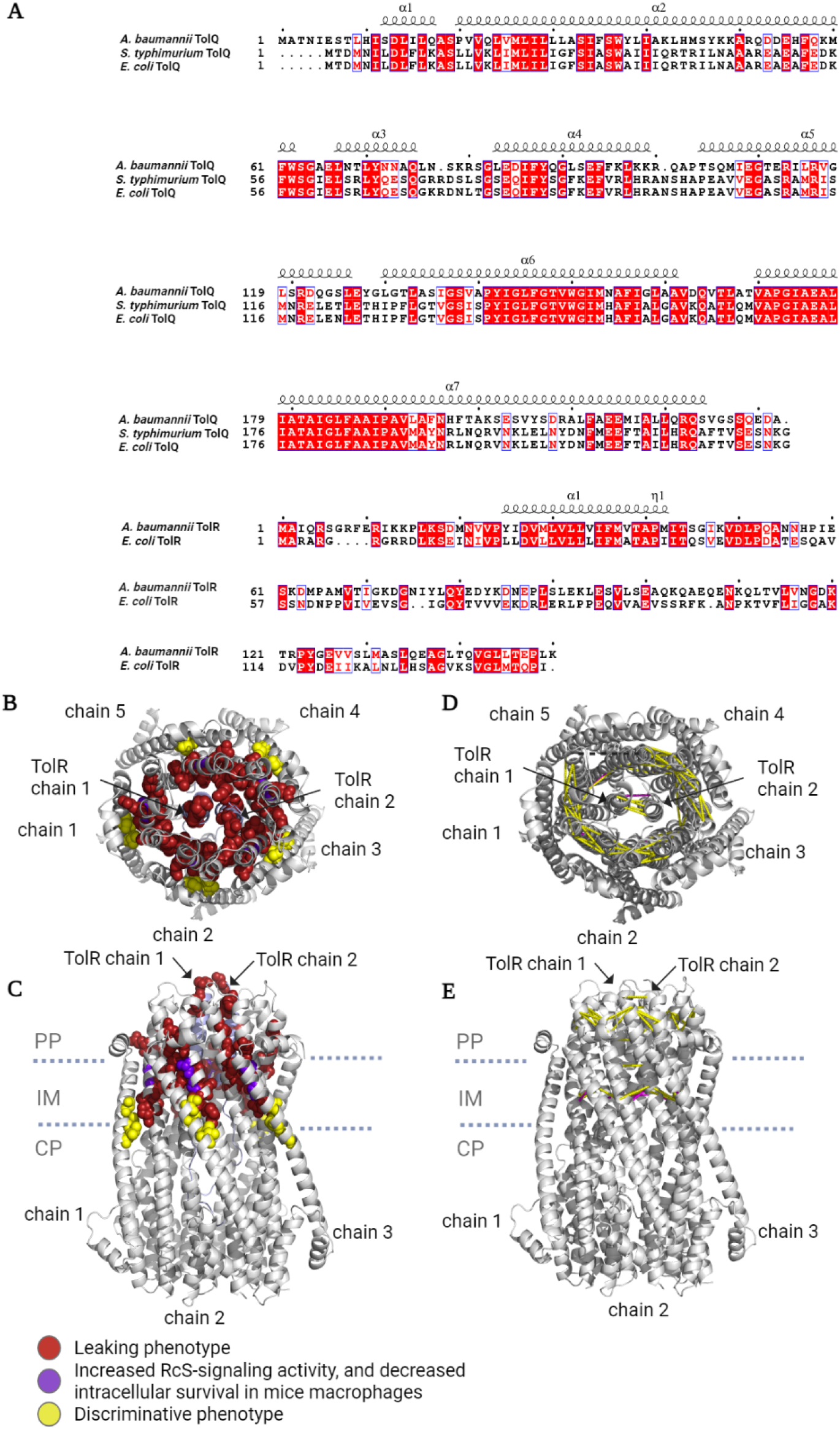
Validation of *Ac*TolQ-TolR structure by prior functional data. **(A)** Sequence alignment of TolQ and TolR in species for which point mutation studies have been conducted. The mutated residues are plotted as spheres on the *Ac*TolQ-TolR structure (gray) on the position of the Cα atoms of homologous residues in *E. coli* and *S. typhimurium* as periplasmic **(B)** and side **(C)** view. **(D, E)** Representation (shown as dotted yellow lines between the homologous Cα atoms) of *E. coli* TolQ and TolR residues that can be crosslinked when they are mutated to cysteines. The dotted purple lines correspond to *in vivo* crosslinking from the prior data and the protomeric non-bonding contacts we observed in *Ac*TolQ5-TolR2 structure.

The single mutations of residues corresponding to Asp^27^ in A. baumanii TolR or Thr^148^ and Thr^181^ residues in *A. baumanii* TolQ result in outer membrane destabilization phenotypically comparable to that observed for *tol* operon deletion in *E. coli* (*23*). This residue colocalizes with Thr^148^ and Thr^181^ in one copy of *Ac*TolQ-TolR at the distance which would allow for transient interactions between these residues (Fig. 3, E and F). A similar arrangement of the corresponding residues has been observed in ExbB-ExbD and MotA-MotB complex structures (*37–39*). Due to the asymmetric position of TolR dimer inside the TolQ pentamer, the Thr^148^ and Thr^181^ residues of each TolQ protomer form multiple interactions with TolR. Specifically, Thr^148^ and Thr^181^ in TolQ chain 1 form non-bonded contacts with Met^29^ and Ile^26^ from TolR chain 1, respectively (fig. S10); Thr^181^ in TolQ chain 5 forms hydrogen bonds and non-polar contacts with Tyr^25^ in TolR chain 2 (fig. S11). In addition to the interactions mentioned above the Asp^27^ from chain 2 of TolR along with Ile^26^ are also within non-bonding contact distances (3.0 - 3.8 Å, respectively) from Leu^145^ residues in the three adjacent TolQ protomers, chains 2, 3, and 4 (Fig. 3G).

Notably, we observed the double conformation of the Phe^146^ located in TMH2 of TolQ chain 1 in the area with the widest distance between TolQ protomers due to asymmetry (Fig. 3H). One possible explanation for the conformational changes interrupting contacts between TolQ protomers could be the requirement of such alterations for TolR entry.

According to the electron density map, the side chain of Phe^146^ either freely points outside the channel or turns towards the TMH2 of the next TolQ protomer chain 5, forming two non-bonding contacts with Leu^185^ (Fig. 3I). The “pointing out” conformation of the Phe^146^ phenyl group was only detected when TolR dimer was present inside TolQ. Similar to Phe^146^ residues in apo TolQ5 (fig. S12), the Phe^146^ residues of the rest four TolQ protomers point towards TMH2 contributing to the interaction interface between TolQ protomers in the *Ac*TolQ5-TolR2 structure (fig. S13).

Individual substitution of several conserved TolQ residues has been shown to significantly alter its cellular function. Specifically, the Pro^190^Val, Glu^176^Gln/Leu, Ala^155^Glu, Gly^144^Ala, Gly^147^Ala, Gly^151^Ala, and Leu^185^Cys mutations in *E. coli* TolQ protein resulted in significant cell envelope defects (*18, 23*). According to *Ac*TolQ5-TolR2 structure, the proline oxygen atom of Pro^190^ forms a main-chain polar contact with the nitrogen atom from Ala^194^ which stabilizes α helical structure of TolQ’s protomers’ TMH3. Substituting Pro^190^ to a Val would extend the distance between the two interacting atoms from 2.9-3.4 to 3.5-3.6 Å, potentially affecting each protomer’s folding in TolQ pentamer. TolQ’s Glu^176^ forms from 1 to 4 main-chain polar contacts stabilizing the helical architecture in TMH3 of each protomer. The Glu^176^Gln substitution either fully abrogates main-chain polar bonds or reduces their number (chains 1, 3, 4, and 5), thus affecting the interactions of this residue with TMH3. Glu^176^Leu substitution would have a milder effect on the main-chain polar bonds. However, the shorter side chain of leucine would fully abrogate interactions of this residue with TMH3. Ala^155^ is involved in main chain polar bonds with Leu^159^ and Gly^151^, which stabilize the position of TMH2. When substituted with Glu, the long negatively charged side chain at this position will promote a significant van der Waals overlap resulting in multiple clashes with residues of TMH3 from the same TolQ protomer (chains 1, 2, and 4 of TolQ). Gly^144^, Gly^147^, and Gly^151^ are located at the pore forming TMH2 of TolQ. Gly^144^ maintains main-chain polar interactions within the same protomer with Thr^148^ and Ala^140^ 2.9-3.1 Å in length. Substitution of Gly^144^ by Ala will result in either abrogation of contact with Ala^140^ and/or elongation of the existing contact distance with Thr^148^ in adjacent protomers. Gly^147^ faces intra-protomeric TMH1 of TolQ and provides polar interactions with Gly^151^ and Ile^143^. The effect of the Ala mutation of this residue mutation may be explained by the loss of conformational flexibility of TMH2 necessary to accommodate the TolR helices. Leu^185^ is located at TMH3 of TolQ and is part of hydrophobic interface between TolQ protomers. A cysteine at this position would introduce a sulfhydryl group abrogating multiple non-polar bonds and thus dramatically affecting the protomer packing of TolQ pentamer.

The substitutions of Ala^180^ or Gly^184^ equivalent residues in *S. typhimurium* TolQ by a Val or an Asp, respectively, increases RcS-signaling activity and suppresses intracellular survival in C57BL/6 mice macrophages during bacteremia (*22*). Gly^184^ is located in TMH3 of TolQ and is proximal to TMH2 of the same protomer. The substitution of Gly^184^ to Asp will cause clashes with TMH2 as well as with TMH1 residues. In turn, the Ala^180^ forms main-chain polar contacts with Ala^177^ and Gly^184^. Mutation of Ala^180^ to Val will introduce multiple clashes with residues from TMH2 from the same protomer.

Cysteine substitutions of Ser^33^, Ser^36^, or Ile^40^ residues in TolQ result in colicin- and DOC- sensitive phenotype (*18*). These three residues are located in the IM region of TMH1 with side chains facing TMH2 or TMH3 of the same TolQ protomer (Fig. 4, B and C). The modeling of Cys at the Ser^33^ position resulted in clashes with residues from TMH3 in each protomer of TolQ. Ser^36^ to Cys mutation will result in clashes with multiple residues from TMH2, TMH1, or TMH3, and abrogation of polar contacts with Ala^32^ and Ser^33^. While the Ile^40^Cys mutation does not produce obvious clashes, its discriminative effect may be explained by its location within Ser^33^-Ala^41^ TolQ region interacting with TolA.

In TolR dimer packed inside TolQ pentamer, the Ala^40^’s side chain faces either TMH3 of the TolQ protomer or towards the chain 1 of TolR, where it forms polar contacts with Phe^36^ and Ile^43^. Mutation of this residue to a Val will cause clashes with TolR’s chain 2 and the residues from TolQ protomer. Pro^41^ residue is located at the extremity of TMD of TolR facing periplasm. Substitution of this residue by a Val introduces multiple clashes with TolQ residues belonging to chains 2 and 3. Pro^24^ is located at the border of the disordered N-terminus region of TolR and well-ordered helical TMD. In TolR chain1, Pro^24^ forms non-bonding contacts with Tyr^145^ of TolQ’s chain1. The Pro^24^Leu mutation will abrogate interaction with Tyr^145^ and will introduce multiple clashes with residues from TolQ chain 2 and Ile^26^ from the TolR’s chain 1. Furthermore, in TolR’s chain 2, this mutation creates multiple clashes with residues from TolQ’s chain 4.

As mentioned above, TolR’s Asp^27^ is expected to play a critical role in the proton pathway. Substituting this negatively charged amino acid to an Ala or a Cys will sabotage such function. Mutation of this residue to glutamic acid will also be detrimental. In TolR chain 1, the Asp^27^Glu mutation introduces multiple clashes with Leu^145^ from TolQ’s chain 2 or Pro^24^ from the same TolR protomer due to its extended side chain.

The analysis of *Ac*TolQ5-TolR2 structure also facilitated the interpretation of previously published *in vivo* crosslinking experiments to map the interface between these proteins in the complex (*17, 18*). Following these previous studies, the residues identified as proximal belong to TolQ helices line at the inner surface of the channel (Fig. 4, D and E). However, the crosslinking observed between residues of chains 1 and 5 of TolQ protomers is not supported by our structural model of *Ac*TolQ-TolR due to the abovementioned asymmetry introduced by the binding of TolR dimer in the complex. It should be noted that the *in vivo* crosslinking studies were conducted for TolR (*17*) and TolQ (*18*) individually. So observed discrepancy may be due to the presence of apo versus TolR-bound TolQ in bacterial cell. The cross-linked pair of residues Tyr^142^-Ile^189^ of TolQ and Val^23^-Val^23^ of TolR correspond to the inter-protomeric non-bonding contacts we observed in *Ac*TolQ5- TolR2 structure (Fig. 4, D and E, and fig. S9).

### Evaluation of *Ac*TolQ5-TolR2 complex structural plasticity and permeability by molecular dynamics

Using the obtained *Ac*TolQ5-TolR2 structure we applied in silico molecular dynamics approach to gain molecular insight into the possible structural fluctuation of this complex in the context of bacterial membrane. The stability of the simulated *Ac*TolQ5 and *Ac*TolQ5-TolR2 structures was maintained throughout the duration of the microsecond simulation timescale. No significant structural rearrangements were observed in either *Ac*TolQ pentamer or *Ac*TolQ-TolR hetero-heptamer in this experimental setup. The final structural models obtained from the simulation studies closely resembled the model derived from cryo-EM studies (Fig. 5A-F). Analysis of the root mean square fluctuations (RMSF) indicated minimal structural fluctuations during the simulation (fig. S14). The TolQ5-TolR2 complex exhibited minor fluctuations in the terminal region, while the TolQ pentamer showed slightly higher fluctuations in the soluble cytoplasmic helix (Leu^67^- Leu^76^) and the adjacent loop region (Trp^62^-Glu^66^) in only one chain. In both cases, the transmembrane portions of the structures retained their shape. The energy landscapes calculated from the simulations were mostly flat, indicating minimal structural rearrangements in the microsecond-long MD simulations. These findings suggest that TolQ5 and TolQ5-TolR2 structures remain stable over extended time periods when analyzed in the context of bacterial membrane mimic and that the molecular model obtained from cryo-EM studies is representative of the native conformation of this Tol-Pal sub-complex.

**Fig. 5.**
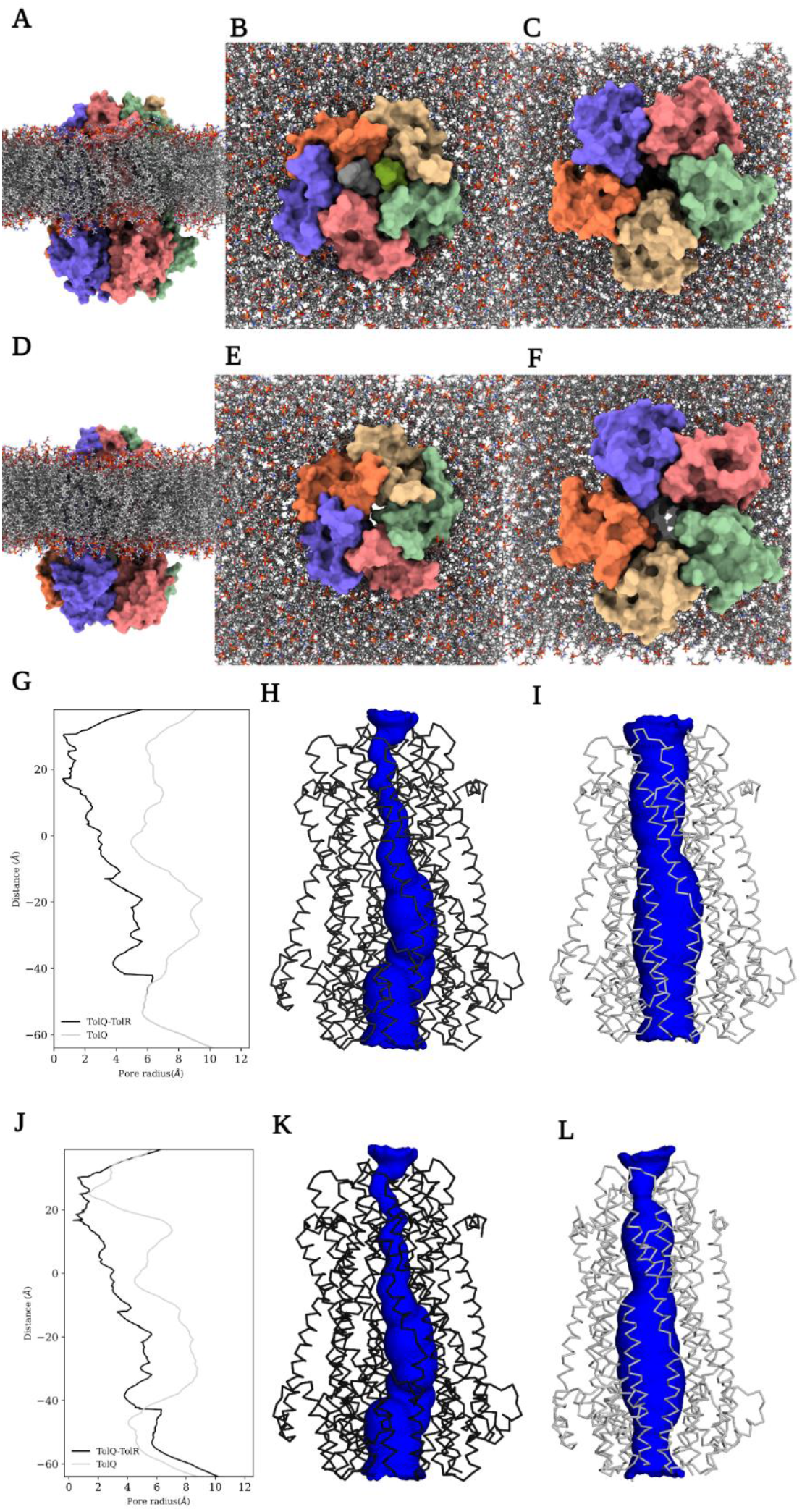
Structural plasticity and pore profiles of *Ac*TolQ-TolR and apo *Ac*TolQ structures. **(A), (B), (C)** TolQ-TolR structure and **(D), (E), (F)** TolQ structure extracted after the MD simulations in native membrane environment. For clarity, only the bilayer is shown. **(A,D)** side view, **(B,E)** periplasmic view, **(C,F)** cytoplasmic view. Protein is shown with surface and the lipids are shown with sticks representations. Both the structures remain largely unperturbed. **(G), (H), (I)** Calculated pore profiles of the cryo-EM structures and **(J), (K), (L)** from average structures from MD simulations. Left panel **(G, J)** shows the variation of the pore along the long-axis (membrane normal) of the proteins. The middle **(H, K)** and the right panels **(I, L)** represent the TolQ-TolR structure and TolQ structure, respectively. Proteins are shown in ribbon and pore profiles are shown in surface.

To investigate the dynamic states of the *Ac*TolQ5-TolR2 complex and whether it could selectively allow the permeation of protons (H+) or small cations like K+, we calculated the pore radius along the obtained cryo-EM structure of the TolQ5-TolR2 complex (Fig. 5G-I). The analysis revealed the presence of a narrow pathway in the channel spanning the transmembrane region, and constrictions were observed between the TolR dimer and chain 2 of TolQ. The pathway alternated between single-file water and constriction points. Three constriction points were found near the TMHs of the TolR dimer. Beyond TolR TMH, the channel path widened to a radius of at least 3 Å and maximum 6 Å in some regions, which would allow for the passage of multiple water molecules or ions. Based on these observations, we suggest that the overall conformation observed in the *Ac*TolQ5- TolR2 complex structure corresponds to a non-permeable state of the channel. To confirm this hypothesis and explore the possibility of attaining a partially permeable state compatible with ion leakage based on the current structure, we performed and analyzed the MD simulations of *Ac*TolQ5-TolR2 complex. The calculated pore profile on an average structure extracted from the MD simulations remained mostly unchanged (Fig. 5J-L), highlighting no significant local rearrangements along the narrowest portion of the channel path. We also examined the possible dynamics of K+ and Cl- ions placed within the channel. The simulation confirmed the possibility of the entrance of such ions from the cytoplasmic side into the large cavity of TolQ pentamer and the possibility of K+ ions translocating along the channel. However, our simulations also suggested that the passage of these ions will be blocked by the constriction points around the upper part of TMHs of the TolR dimer (fig. S15). These simulations also suggested that no chloride ions would be favored to enter the channel, as indicated by its electronegative nature. In summary, our simulation found no possible route for uninterrupted ion transfer in the obtained structure of TolQ5-TolR2 complex. To enable the passage of water and protons/ions, significant structural rearrangement of the TolR dimer in relation to the TolQ pentamer will be necessary to allow for passage through identified constriction points. Alternatively, the passage would have to involve other structural states permissive of ion transfer.

We also conducted simulations to check the permeability of TolQ5 in its apo form. Initially, we hypothesized that TolQ5 would lack stability and selective permeability by protons (H+) or potassium ions (K+) due to the larger radius of its pore compared to canonical pores found in other ion-channels/transporters (*48*). In the absence of TolR2, the radius of in TolQ5 channel varies from 3.16 Å at its most narrow point to the maximum of ∼11 Å at the certain segment and having a radius of ∼4 Å in the periplasmic side (Fig. 5G- I). To explore the dynamics of unbound TolQ5 and assess this protein’s potential for nonselective ion permeation, we conducted the following MD calculations.

During the simulations, the channel in TolQ5 remained continuous thus confirming the possibility of this protein forming a pore in the IM. The narrowest region of the pore with a radius of 3.16 Å was at the cytoplasmic side and ∼4 Å at the periplasmic side with Leu^167^ residues from all five TolQ protomers forming a constriction point. The simulations suggested rapid dewetting in specific regions of TolQ5 pore, and these areas remained water molecule-free throughout the simulation, even after simulating with generated pore waters (fig. S16). Dewetting initiated in the lower part of the channel, where the radius was approximately 3.16 Å, while the dehydrated portion of the channel in other regions with pore radius reaching 7.24 Å was considerably larger. Parts of the two TMHs formed the dewetting pore from each TolQ5 protomer (-^167^LATVAPGIAEALIATAIGL^185^- from TMH3 and -^144^GLFGTVWGIMNAFIGLAAV^162^- from TMH2). The formation of these extensively dewetted regions would not permit the continuous passage of K+/H+ ions through TolQ5 (fig. S17).

Based on the simulations, the structures appear to have a stable conformation that corresponds to the non-permeable state of the AcTolQ5-TolR2 stator unit, and due to the highly ordered plugging TMHs of TolR into the TolQ pore, ions are inhibited from diffusing freely across the pore.

### Analysis of AcTolQ-TolR complex dynamics

Next, we explored the dynamics of component interactions in *Ac*TolQ5-TolR2 complex by analyzing the heterogeneity of the particles used in the cryo-EM 3D reconstruction through 3D Variability Analysis (3DVA) (*49*). The 3DVA algorithm identifies the ‘variability components’ of the particles in the latent space, which may be indicative of different conformational states captured in the analyzed protein sample.

Accordingly, this analysis identified multiple types of motions within the *Ac*TolQ5-TolR2. Notably, we observed that the shape of the TolQ5 pore undergoes changes primarily driven by the bending motions of its protomers (SM 1, cytoplasmic and side view).

Furthermore, the rotation of a specific TolQ protomer around the perpendicular axis within the cytoplasmic region played a role in modulating pore shape in several variability components as observed in the cytoplasm and cross view of Video 2 and 3 (SM, 2 and 3).

We observed only marginal movements in the central and upper segments of the TolR TMHs, which provided additional support for a stable H+ non-permeable state of the stator unit structure. In contrast, the 3DVA analysis suggested potential dynamic folding and refolding processes occurring at the N-terminus of TolR in the non-permeable state of the *Ac*TolQ5-TolR2 complex. The density corresponding to the disordered N-terminus of TolR and the N-terminal part TMH altered between defined or undefined across all five analyzed modes (SM1-5, sliced view). This recurring pattern can be indicative of this portion of TolR being in dynamic equilibrium between ordered and disordered states or accompanied and involved in the movement up and down the TolQ cavity which results in averaging out the molecular density.

Taken together, 3DVA analysis revealed the bending and rotation of TolQ protomers with the magnitude of TMH movements smaller than those occurring in the cytoplasmic region. These coordinated structural changes of TolQ protomers, along with their collective interactions, have the potential to modify the shape of the pore/channel and exert an influence on its interaction with TolR dimer components.

### Modeling of TolQ-TolR interactions with TolA

Previous analysis (*1, 21, 50*) suggested direct interactions between TolQ-TolR sub- complex with the N-terminal portion of TolA protein localized to IM. Therefore, we used the obtained structure of *Ac*TolQ-TolR to model its interaction with TolA using AF2 ColabPro server (*51*).

In the highest-ranked predicted model of TolQ-TolR-TolA complex, the single TMH of TolA spanning TM residues Phe^9^ to Leu^33^ is oriented at ABC angle with respect to the first TMH of TolQ and the interactions between the two structural elements are stabilized by van der Waals interactions close to the cytoplasmic side of the complex (Fig. 6, A and B). Recently published Rosetta model predicted that *E. coli* TolA TMH is inserted into a TolQ groove on the exterior of the TolQ pentamer (*52*). However, cryo-EM reconstruction of the ExbB-ExbD-TonB system in *Pseudomonas savastanoi* (*Ps*ExbB-ExbD-TonB), which share significant sequence similarity with Tol-Pal system components revealed an arrangement similar to the one suggested by our model (Fig. 6C). Analysis of *Ps*ExbB- ExbD-TonB data at a low contour level identified a single additional density, attributed to TMH of TonB, a TolA homolog, traversing the micelle located on the outer side of the ExbB-ExbD complex. This density is positioned to interact with ExbB, a homolog of TolQ, aligning with the predicted location of TonB, a homolog of TolA, determined by covariance analysis (*39*). The same arrangement between TolA and TolQ is suggested by our AlphaFold model. According to this model, residues Ala^15^, Thr^19^, His^23^, and Leu^30^ belonging to TolA’s TMH are involved in non-bonding contacts with Ser^33^, Ile^34^, Trp^37^, Tyr^38^, and Ala^41^ residues of the TolQ TMH1. Notably, these residues are conserved across the corresponding components of *Ac*Tol-Pal and *Ps*ExbB-ExbD-TonB systems, thus further validating the proposed model and suggesting that TolQ-TolA interactions may be following the same pattern as observed in ExbB-TonB complex structure (*39*). The polar interaction between *Ac*TolA His^23^ and *Ac*TolQ Ser^33^ was also suggested by our model, which is consistent with previous observations in *E. coli* TolA-TolQ and *P. savastanoi* TonB-ExbB systems (*39, 52*).

**Fig. 6.**
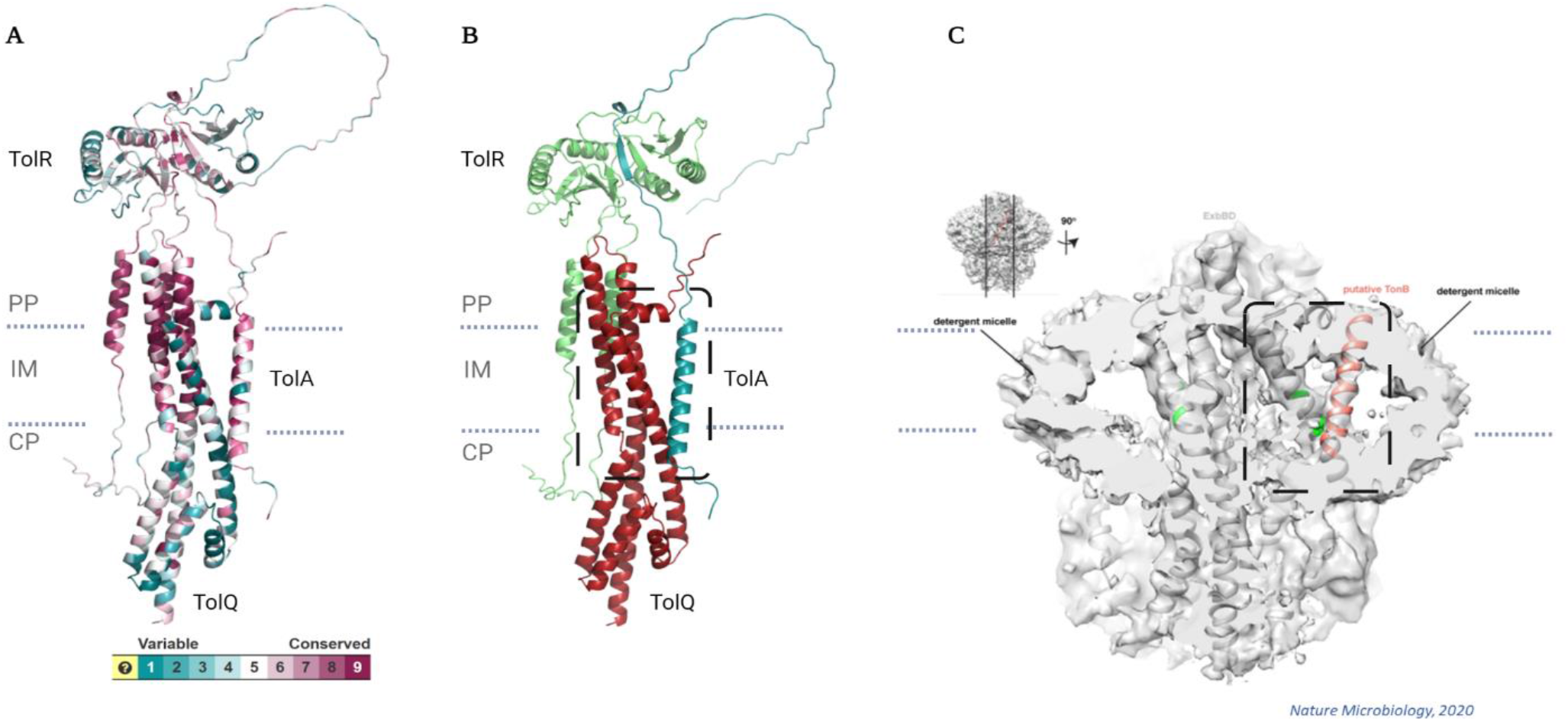
Interaction of *Ac*TolQ-TolR with TolA. **(A)** AlphaFold2 model colored by residue conservation. **(B)** AlphaFold2 model colored by subunit (TolQ-red, TolR- green, TolA-blue). **(C)** Low-resolution cryo-EM map of ExbB-ExbD-TonB adapted from (*39*).

Our *Ac*TolQ-TolR-TolA model suggested that TolA is forming direct interactions with the periplasmic domain of TolQ. More specifically, TolA sequence following the TMH, which makes part of its periplasmic extended domain, is inserted into the C-terminus periplasmic domain of TolR to form two parallel β-strands (Fig. 6, A and B). This arrangement also positions the five residues of TolA (Thr^46^-Val^50^) between protomers in TolR dimers proximal enough to form multiple hydrogen bonds and non-bonded contacts with TolR residues. This model is somewhat reminiscent of interactions observed in recently defined crystal structures of *E. coli* ExbD-TonB periplasmic fragment complex, where ExbD, the TolQ homolog, captures one copy of the TonB peptide via β-strand recruitment (*53, 54*). However, the superimposition of this crystal structure with the obtained model of *Ac*TolQ-TolR-TolA highlights a significant difference in the orientation of TonB versus TolA interacting regions. Thus, the experimental validation of proposed *Ac*TolQ-TolA interactions is required.

## Discussion

The components of the Tol-Pal system (TolA, TolB, TolQ, TolR, and Pal) assemble into complexes that, depending on their composition, are anchored at the inner and/or outer membranes and interact with peptidoglycan. The structural transitions of these complexes are driven by the electrochemical gradient that forms between the cytoplasm and periplasm. The TolQ-TolR-TolA subcomplex is considered a motor that generates rotary force driven by the electrochemical gradient. The rotary force is then transferred by TolA to reorganize other subcomplexes of the Tol-Pal system. There are unanswered questions about how TolQ-TolR generates torque and how the torque force is used by TolA. The torque generation is presumably conserved across other reported 5:2 motors; however, whether the force is used in an analogous way after it is generated remains to be determined.

The structures of the pentameric apo TolQ pentamer and the TolQ-TolR 5:2 complex are used as the basis for the TolQ-TolR-TolA activation mechanism proposed here, which has distinct features not considered in previously published models. The earlier models were based on the similarity of the TolQ-TolR system to two other systems: the MotA-MotB pilus motor and the ExbB-ExbD transport system. However, the structures we present have important features that differentiate TolQ-TolR from the structures of homologous systems: (1) TolR helices in the TolQ pore are related to each other by translation, not by rotation as in the homologous structures of MotA-MotB and ExbB-ExbD complexes; (2) the TolQ pentamer in the 5:2 complex deviates from perfect 5-fold symmetry in periplasmic region much more than in other complexes so that contacts between two subunits of TolQ are broken (chain 1/A and chain 5/E in our model).

In our model, the TolQ-TolR-TolA complex is assembled as follows. TolQ is embedded in the inner membrane (IM), with two N-terminal fragments of two TolR molecules complexed inside the pentameric TolQ pore. The periplasmic domains of TolR form a dimer and are flexibly attached to the helical portion of TolR complexed with TolQ. The N-terminal transmembrane helix of TolA complexes with the TolQ-TolR subcomplex, and its periplasmic part can also interact with TolR. TolA is also thought to extend through the periplasm and is known to interact with TolB and Pal through its C-terminal domain in a PMF-dependent manner. This assembly is consistent with other biochemical, biophysical and phenotypical data.

In this assembly, The TolQ-TolR complex forms a stator-rotor structure in the inner membrane. The stator-rotor molecular motors operate by cycling through different conformational states, typically three, in response to an energy source. During operation, one component of the motor rotates relative to the other. The number of cycles involved in a full 360° rotation of the stator-rotor motor depends on the intermediate conformational states. In the MotA-MotB system, ten cycles were assumed, each at 36°. However, a model with a 72° motor cycle has also been discussed as a possibility (*38*).

Based on our structure, we propose a model for the TolQ-TolR system that reconciles the translational symmetry between the TolR molecules, a feature not previously discussed. We propose that the transfer of one proton generates a conformational change in the TolQ-TolR complex, resulting in a 36° rotation of TolQ, coordinated with a 180° rotation of the TolR helices within the TolQ pore. This new structural state recreates the same interactions between TolQ and TolR as the starting point but the positions of two TolR helices are swapped. Thus, the entire complex requires another 36° rotation coupled to another proton transfer to return to the state with the same stereochemistry of TolQ and TolR interactions. Since the local chemical interactions between the TolQ and TolR helices are maintained after the first 36° rotation, the kinetics of the second step is the same as in the first step. Ultimately, in our model, the entire TolQ-TolR cycle involves a 72° rotation of the TolQ pentamer around the TolR dimer, forming a state shifted by one TolQ subunit (Fig. 7). In each 36° cycle, when the two screws rotate in opposite directions, their contacts are preserved. This principle underlies the action of a twin-screw pump. Therefore, due to the helices’ screw symmetry, the simplest solution for achieving a 180° rotation is for the TolR helices to rotate in opposite directions. In such a model, the distinction between stator and rotor is difficult because both components (TolQ and TolR) undergo significant structural changes. In the MotA-MotB and ExbB-ExbD systems, such rearrangements were not proposed because their inner components were related by twofold symmetry, recreating equivalent structures after a 36° rotation.

**Fig. 7.**
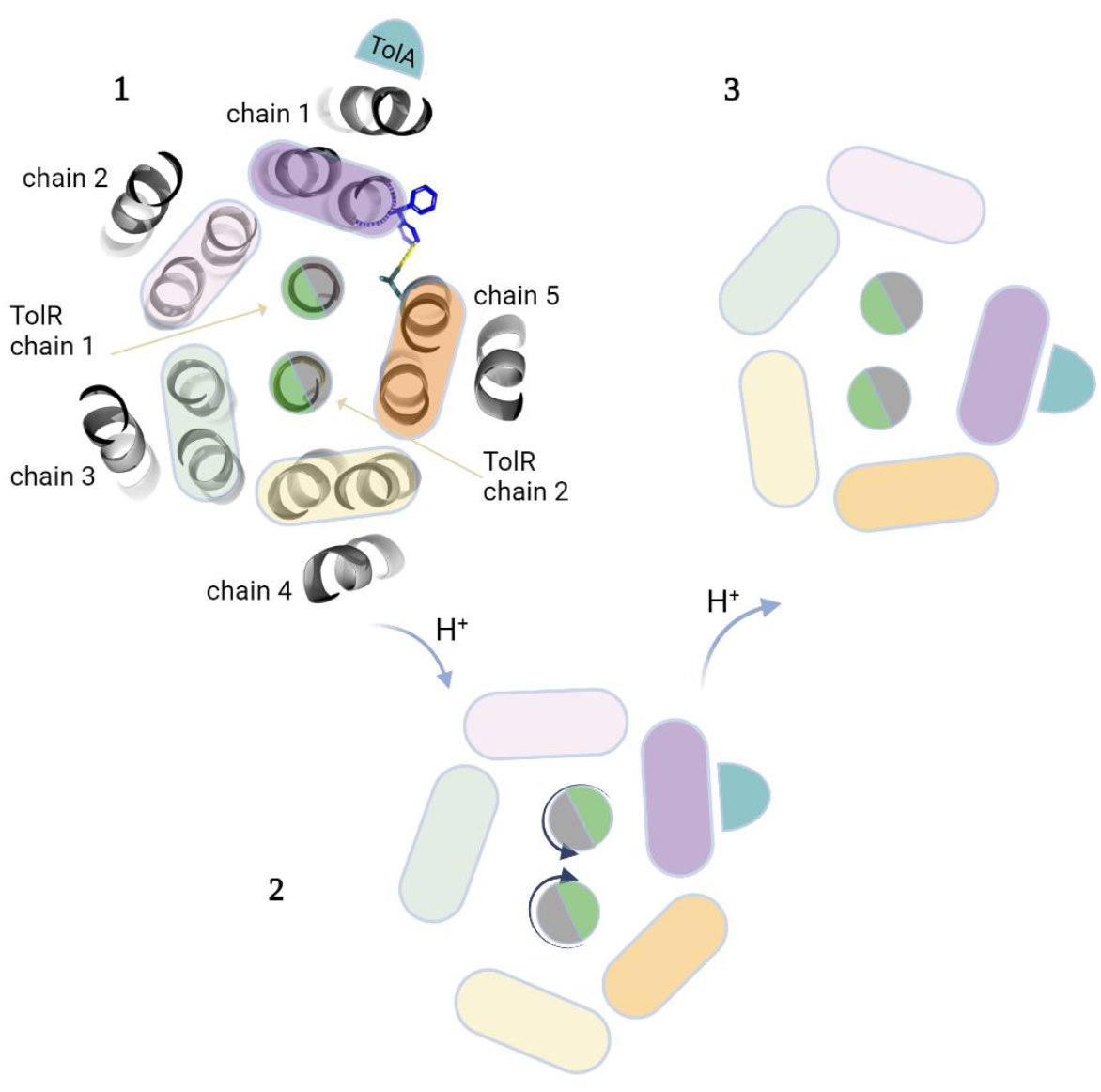
Mechanistic model of action with TMD rotational movements of *Ac*TolQ and *Ac*TolR protomers in IM environment. State 1 represents a cartoon showing the TMH core of the *Ac*TolQ-TolR complex, viewed from the periplasmic side, with five TolQ chains asymmetrically surrounding two TolR dimer TMHs and TolA single TMH bond to TolQ TMH chain 1/A protomer. The actual view of *Ac*TolQ- TolR complex from the periplasmic side is overlapped for convenience. State 2 represents the same complex with 36° rotation of TolQ pentamer around TolR dimer after 180° rotation of each TolR protomer. State 3 represents the same complex with 72° rotation of TolQ pentamer around TolR dimer.

The rotation of TolQ-TolR in the IM must be coupled with changes in other components of the Tol-Pal system in the IM, periplasm, and OM. For a torque-generating motor, energy propagation requires that both components be anchored separately, meaning that both TolQ and TolR must be externally anchored. TolA is the protein that anchors TolQ to a structure external to TolQ. One postulated possibility is TolA’s interaction with TolB, which is anchored in the OM by Pal; such interactions would create a necessary anchor for TolQ. In this model, TolR is anchored through interactions with peptidoglycan. However, there is also an alternative solution (*55*), in which TolA is topologically trapped by the TolR dimer, simultaneously creating both anchors, with the rotation of TolQ around TolR pulling TolA through the topological trap. Our modeling with AF2 and the recently published TolR homolog ExbD structures complexed with a TolA homolog TonB and HasB (TonB paralog with single heme transport function) peptides (*53, 54*) could correspond to intermediate structures leading to the formation of such a topological trap.

Collectively, experimental data and our analysis indicate that both solutions are likely used, probably for separate functions, or simultaneously.

As discussed above, our analysis suggests that the obtained structure of *Ac*TolQ5-TolR2 complex corresponds to a non-permeable conformation state (Fig. 8A). TMHs of the TolR dimer are highly ordered and embedded inside the TolQ pentamer forming a transmembrane IM channel with no ion flow. The periplasmic domain of TolR seems to be dimerized in this state as well. The periplasmic and partial IM region of one TolQ protomer is shifted to the side due to the insertion of the two TolR TMHs. Considering that PG is expected to be rigid (*56*) and that the periplasmic domain of TolR needs to bind PG for its active state (*12*), we propose a rotary mechanistic model for the TolQ-TolR complex where the TolQ pentamer rotation around TolR dimer during activated phase, similar to the ExbB-ExbD mechanistic rotary model (Fig. 8, B and C). The activation of the proton flow can be initialized by the Thr^46^-Val^50^ sequence of TolA embedding into the β-sheets predominantly formed by the C-terminus of the TolR dimer. In this case, the TolA hairpin may need no PG pore to extend to OM to associate with TolB in its complex with Pal (Fig. 8B). The initialization state of the activated TolQ-TolR-TolA complex may be supplemented with simultaneous 36° rotation of TolQ pentamer and 180° rotations of TolR protomers discussed above to fit TolA in a periplasmic domain. Deprotonation of stator residues orchestrated with another 36° rotation of TolQ (72° in sum) drives relaxation of the TolA helical hairpin toward the IM (Fig. 8C). We did not indicate the elongation of the flexible TolR linker between TMHs and the beta-sheet periplasmic domain during the activation step because the thickness of periplasm varies between bacterial species (*57*). The distance between the IM and the low-resolution density of the TolR periplasmic domain in the low contour cryo-EM map is about 50 Å, (fig. S7) which seems to be enough to reach and go through the thin PG layer.

**Fig. 8.**
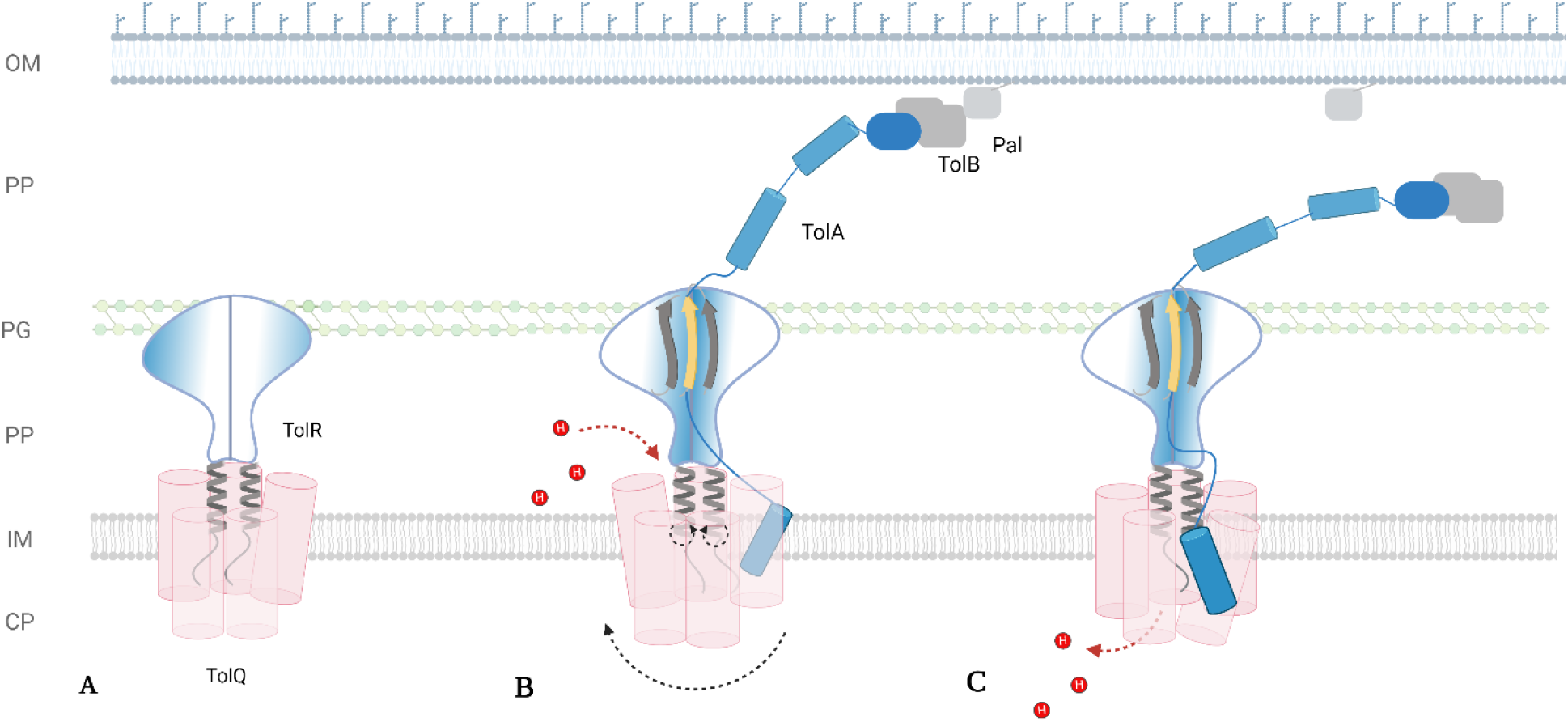
Mechanistic model for the generation of Tol-Pal torque. **(A)** Structurally resolved non-permeable state of asymmetric *Ac*TolQ-TolR complex. **(B) ‘**Activated’ state of *Ac*TolQ-TolR-TolA complex depicting asymmetric TolQ pentamer rotated 36° around TolR dimer which in turn finished 180°- rotation creating at periplasmic region β-sheet with TolA β-strand. This allows TolA to bind TolB-Pal OM associated sub-complex **(C)** Deprotonation state supplemented with the next 36° rotation of asymmetric TolQ pentamer with relaxation of the TolA helical hairpin toward the IM.

The structure of the TolQ-TolR system from *E. coli*, recently published (41), should be cautiously interpreted when compared to our results. The molecular density maps obtained in that study have low resolution (4.7 Å for TolQ-TolR complex). As a result, accurately assigning residues’ positions in the density map was not possible, and the model was obtained by docking the AlphaFold model directly into the density. This atomic model does not agree well with the density maps deposited (EMDB-16816). Upon inspection, we found that out of the five subunits of *Ec*TolQ in the AlphaFold model, only three partially agree with the map, while the remaining two subunits do not. The density in these regions indicates that the reconstruction contains a mixture of structural states. This interpretation is further supported by the presence of molecular density representing a single helix that is not discussed but overlaps with both poorly resolved subunits of the AlphaFold model simultaneously.

In conclusion, here we reported the apo TolQ and TolQ-TolR complex structures of *A. baumannii*, a pathogen of significant clinical concern. We analyzed computationally the molecular flexibility and characterized the proton sites and general flow path through the structures. Our TolQ-TolR structure is in a state that does not allow for proton flow. The phenomenon of dewetting regions of the pore formed by TolQ protomers identified through molecular simulations holds significant scientific interest and may serve as inspiration for the design of de-novo hydrophobic pores. We mapped critical functional mutations that disrupt the bacterial OM stability onto our TolQ-TolR structure identifying regions that could be used in rational, structure-assisted development of antibacterial therapeutics. We predicted complexes of TolQ-TolR-TolA and discussed possible mechanisms of actions of TolQ-TolR-TolA subcomplex of Tol-Pal, which need to be experimentally validated by further structural studies involving co-expression of all three IM protein members. Our findings bring new knowledge to help further understanding of the mechanism of how molecular stator units energize vital bacterial functions, which will potentially be applicable for antibiotic drug discovery.

## Materials and Methods

### Protein cloning, expression, and purification

The *A. baumannii* DNA encoding the protein *Ac*TolQ (GenBank: KFC02860.1) was codon optimized for expression in E. coli expression systems, synthesized (Twist Bioscience), and used as the template for subcloning into pMCSG53 and pNIC-CH vectors with an amino- and carboxy- terminals 6His-tag, respectively. Both constructs possessed high level of expression. Therefore, we proceeded further with the construct cloned in pMCSG53 since it contained a TEV cleavage site enabling us to eliminate all non-specific Ni resin binders from the samples. For the *Ac*TolQ-TolR complex, the *Ac*TolR (GenBank: HAV5244423.1) sequence was also codon-optimized, synthesized (Bio Basic Inc.), and used for subcloning. Initially, we utilized a single-cassette- polycistronic strategy (*58*) using the mentioned above pMCSG53 vector where 6His-tag was associated with *Ac*TolQ protein. This led to higher abundance of *Ac*TolQ than *Ac*TolR in the purified sample. To overcome these technical difficulties, we applied a multi-cassette approach (*58*) where N-terminally 6His-TEVcleavage site-tagged *Ac*TolR (multiple cloning site 1 (MCS1)) and untagged *Ac*TolQ (MCS2) genes were cloned in the petDUET1 vector. Vectors were transformed into the C43(DE3) competent cells by 60- second heat shock at 42°C, and 1h outgrowth, and plated onto LB agar selection plates.

Subsequently, the single colonies of transformed cells were cultured overnight in 50 ml LB media supplemented with ampicillin (100 µg/ml) at 37°C. Next morning, the 3, 5, or 6 L of LB media containing ampicillin (100 µg/ml) were inoculated with starter culture for further growth at 37°C until reaching an OD600 of 0.6. Afterward, the culture was incubated on ice for 1 h, followed by induction using 1 mM isopropyl beta-D-thiogalactopyranoside (IPTG), and overnight growth at 20°C. Harvesting of cells was achieved through centrifugation, and pelleted cells were resuspended in 1xPBS (pH = 7.4) and lysed using serial freezing and thawing cycles in the presence of phenylmethylsulphonyl fluoride (PMSF), DNase1 (20 ug/ml), Lysozyme (100 ug/ml), and 0.5 mM reducing agent (TCEP). All subsequent purification steps were carried out at 4°C. Cells were broken in a cell disruptor (Avastin) at 15,000psi, 3 passes. Cell debris was spun down at 13,500xg for 20 minutes. The isolated supernatant was ultracentrifuged for 1h 40min at 40,000 rpm. The pellet (crude membrane preparation) was washed using a spatula with 1xPBS with 0.5 mM TCEP and ultracentrifuged again at 40,000 rpm for 1h 30min. The isolated membrane pellet was frozen and kept at −80°C. The crude membrane preparation was solubilized with extraction buffer (50mM Na Phosphate buffer pH=7.6, 300 mM NaCl, 20% (v/v) Glycerol, 0.5 mM TCEP, 1% DDM) for 2 h. After ultracentrifugation (40 min 40,000 rpm), the supernatant was incubated with 3 ml nickel-nitrilotriacetic acid Ni-NTA beads (Qiagen) preequilibrated with the same buffer overnight. Next day, the beads were washed with 30 ml of wash 1 (50 mM Na Phosphate pH = 7.6, 500 mM NaCl, 15% (v/v) Glycerol, 0.5 mM TCEP, 40 mM Imidazole, 0.1% (w/v) DDM) and wash 2 (50 mM Na Phosphate pH = 7.6, 300 mM NaCl, 10% (v/v) Glycerol, 0.5 mM TCEP, 50 mM Imidazole, 0.05% (w/v) DDM) buffers and then eluted with 15 ml of the elution buffer (50 mM Na Phosphate pH = 7.6, 300 mM NaCl, 5% (v/v) Glycerol, 0.5 mM TCEP, 0.05% (w/v) DDM, 300 mM Imidazole). The eluate was dialyzed overnight with TEV protease (purified in-house) in dialysis buffer (50 mM Tris pH = 8.0, 300 mM NaCl, 5% (v/v) Glycerol, 0.5 mM TCEP, 0.05% (w/v) DDM). The next day IMAC Ni-NTA beads (Qiagen) were used for the second time where the flow-through fraction was passed for 5 times through the beads. The further steps of purification using various detergent environments failed to generate samples suitable for cryo-EM single particle reconstruction. Therefore, other membrane protein stabilizers were tested. Amphipol A8-35 (Anatrace) resulted in the best-quality sample in our trials and was then used in large-scale purifications. Amphipol was added to the flow-through solution in 1/3 - w/w - protein/amphipol ratio and the mixture was incubated with gentle rotation. After 4 hours of rotation, Bio-Beads (Biorad) were added to the mixture in a ratio of 15 mg beads per 1 ml mixture volume and incubated overnight. The next day, the sample with proteins stabilized in amphipol was poured over an empty column to get rid of Bio-Beads and concentrated using 100 kDa cut-off centrifugal concentration tube (Amicon). The concentrated sample was loaded on a Superdex 200 increase 10/300 column equilibrated with detergent-free 50 mM Tris pH = 8.0, 150 mM NaCl, 0.5 mM TCEP Cryo-EM buffer. The peak fractions were collected, concentrated, aliquoted at 15 µl and flash frozen in liquid nitrogen and kept frozen at −80 °C until use. Mass photometry assay was conducted as described previously (*59*).

### Grids preparation and data collection

The purified proteins samples were processed on Quantifoil R 1.2/1.3 300 Mesh Gold grids, which had been made hydrophilic through glow discharge for 90 seconds at 30 mA in a PELCO easiGlow™ system. These prepared grids were then used to create vitrified samples using a Thermo Scientific Vitrobot Mark IV, applying 3 μL of protein at 4°C and 100% humidity before blotting. Data collection was performed using a Titan Krios G2 microscope with a K3 Summit camera, without a phase plate or objective aperture, utilizing SerialEM for automated collection and a GIF Quantum Energy Filter. Data was collected using a 300 kV Titan Krios G2 microscope equipped with a K3 Summit camera, without a phase plate or objective aperture, at a magnification of 105,000× and a pixel size of 0.834 Å. SerialEM software facilitated automated collection in beam-image shift mode, capturing nine images per stage shift across a defocus range of -1.0 to -3.0 μm, with beam-image shift compensation. The slit width of the GIF Quantum Energy Filter was set to 25 eV. Movies were dose-fractionated into 125 frames with a total dose of ∼100 e-/Å2.

### Cryo-EM image processing and model building

For apo *Ac*TolQ, 2,763 movies were imported into CryoSPARC, followed by patch motion correction using binning of 2 and patch CTF correction. Particles were first picked manually to generate templates, which was followed by template-based autopicking, resulting in 472,724 particles. Iterative 2D classification led to the selection of 246,741 particles for further processing. *Ab initio* reconstruction on the clean particle stack with four classes and one round of heterogeneous refinement was conducted, resulting in one class corresponding to the TolQ pentamer and three classes unrelated to TolQ. Non-uniform refinement of 215,170 particles resulted in a 3.02 Å final map in C5 symmetry, which was post-processed using DeepEMHancer to guide map interpretation. The homology model built in SwissModel using the ExbB structure as the template was used in Chimera to build the pentamer. The pentameric model was manually adjusted with rigid body refinement in Coot and then further adjusted using real-space fitting procedures. The built model was refined in Servalcat within CCPEM and validated with the MolProbity server. Model statistics can be found in table S2.

For *Ac*TolQ-TolR complex, all movies were imported into CryoSPARC followed by patch motion correction using binning of 2 and patch CTF correction (*60*). Particles were first picked using blob picker on 200 micrographs to generate initial 2D templates. Templates were created from selected 2D classes and template picking on all micrographs, resulting in a total of 3,009,251 particles. Particles were extracted with box size of 432 pixels followed by several rounds of 2D classification that yielded 465,839 clean particles. *Ab initio* reconstruction on the clean particle stack with three classes to separate TolQ/TolR complexes from TolQ oligomers and contaminants. The class for TolQ/TolR complex (224,768 particles) was selected for homogeneous refinement and non-uniform refinement in C1 symmetry and resulted in a resolution of 3.46 Å (*61*). The periplasmic domain of TolR was not well resolved due to its small size and high flexibility and was likely causing misalignment of the well-resolved transmembrane domain. For this reason, local refinement was performed by masking out the density from the periplasmic domain, which improved the map resolution to 3.34 Å. An initial atomic model was obtained by manually docking the AF2 model of the TolQ-TolR complex to the map using Chimera (*62*). The model was then iteratively rebuilt manually in Coot and refined using *Phenix.real_space_refine* and Servalcat (*63–68*) . Model statistics can be found in table S3.

### Point mutation analysis

The mutagenesis wizard of PyMOL was utilized to analyze mutations mapped onto the TolQ-TolR complex structure (69).

### Molecular dynamic simulation procedure

All atom MD simulations (table S4) were performed for both the TolQ-TolR 5:2 complex and TolQ pentamer. Both the complexes were embedded in a bacterial inner membrane mimic. The membrane model consists of PPPE, PVPG, PVCL2 and POPA lipids in the following ratio PE:PG:CL:PA = 78:12:6:4; and the model is consistent with other theoretical models and experimental assays in the literature (*12, 70–72*). Lipids were distributed equally in both leaflets. A total of 400 lipids were used for both the systems. A water buffer of 14.5 Å is used in z-dimension, with a salt concentration of 150 mM KCl. For the TolQ system, we generated pore water in the empty space of TolR dimers. The systems were generated using Charmm-GUI membrane builder (*73–75*). All the simulations were performed using NAMD (v 2.14) (*76*). We used CHARMM36 force field parameters for these simulations (*77, 78*). The systems were initially minimized for 10000 steps with a conjugate gradient algorithm before the six-step equilibration protocol recommended by Charmm-GUI. Later, we carried out 1 μs simulation for both systems.

The simulations were carried out at 310 K. A cut-off distance of 12 Å was used and the switch distance was set to 10 Å. Long range electrostatics calculations beyond the cut-off were taken into account using Particle Mesh Ewald (PME) method (*79*). A time step of 2 fs was used for the production simulations. The temperature was controlled using the Langevin dynamics (temperature damping coefficient: 1.0) and the pressure was controlled using the Langevin piston method (*79*) (set to 1 atm, oscillation period: 50 fs, damping time scale: 25 fs). Semi-isotropic pressure coupling was used to retain the bilayer shape. Analyses were carried out using VMD (*80*), MDAnalysis (*81*), HOLE (*82*).

### 3D variability assay procedure

To visualize the motion of the complex, a mask that excludes the amphipol ring around the transmembrane domain was generated. This mask and the particles in the local refinement were used as inputs for the 3D Variability job in CryoSPARC (*49*). Five modes were solved in the analysis with C1 symmetry and filter resolution of 6 Å. Results were exported using 3D Variability Display job (simple mode) with 20 frames per mode.

Movies for each mode/component were generated by Chimera (62).

### AlphaFold modeling

To solve the Cryo-EM structures, the AlphaFold models of apo *Ac*TolQ5 and *Ac*TolQ5- TolR2 were calculated using the Google Colab platform and AlphaFold2_advanced option https://colab.research.google.com/github/sokrypton/ColabFold/blob/main/beta/AlphaFold 2_advanced.ipynb (*83*). The default mode of sampling options was used: num_models = 5, ptm option, num_ensemble = 1, max_cycles = 3, num_relax=0, tol=0, num_samples = 1. The model with the highest-ranked pTMscore was used as a template for molecular replacement. For *Ac*TolQ5-TolR2 hetero-heptamer prediction, the 5:2 stoichiometry was included in the input. To stay within the 1400 residues limit, each TolQ protomer was 12 and 9 amino acids truncated from N and C termini, respectively.

The same approach was used to model *Ac*TolQ-TolR interaction with *Ac*TolA. To not exceed the residue limit, TolQ:TolA:TolR in 1:1:2 stoichiometry was used as input.

*Ac*TolQ had no truncations in this calculation set. *Ac*TolA fragment contained the first 99 amino acids corresponding to the inner-membrane domain and the periplasmic fragment. *Ac*TolR dimer was included in full length. The structure figures were drawn with PYMOL (*69*).

## Figures preparation

Figures were generated using Chimera (*62*), ChimeraX (*84*), PyMOL (*69*), VMD (*80*), HOLE (*82*), Matplotlib (*85*), PDBSum (*86*), ENDscript (*87*), PISA (*87*), and BioRender.com.

## Acknowledgments

We thank our colleagues Cameron Samper (UofC) for his valuable suggestions in cloning AcTolQ-TolR construct, Nobuhiko Watanabe (UofC) and Peter Stogios (UofT) for discussion of protein purification.

## Author contributions

Conceptualization: EK, YG, HMK, PT, DB, ZO and AS conceived the project. Methodology: AS, EK, YG, HMK, RDL, BQ, and TE designed the experiments. Investigation: RDL cloned AcTolQ construct; EK cloned AcTolQ-TolR constructs; EK performed expression and purification of all constructs; EK performed screening to stabilize all purified membrane proteins and mass photometry assays; TE performed mass photometry assays and grids preparation; BQ and YG performed single-particle reconstruction; YG, DB, ZO validated models and maps; YG performed 3D visualization assay; HMK performed MD simulations; EK performed structure guided analysis and AlphaFold2 modeling; EK, DB, ZO worked on mechanistic models.

Supervision: AS, DB, ZO, PT. Writing - original draft: EK.

Writing - review & editing: EK, YG, HMK, ZO, PT, DB, AS.

## Data and materials availability

The atomic coordinates of apo *Ac*TolQ structure (PDB: 9AVI) and maps (EMD-43902) have been deposited in the Protein Data Bank (http://wwpdb.org/) and the Electron Microscopy Data Bank (https://www.ebi.ac.uk/emdb/), respectively. The *Ac*TolQ-TolR structure (PDB:8VLW) and maps (EMD:43346) can be found in the mentioned above databases as well.

## References

1. E. Cascales, R. Lloubès, J. N. Sturgis, The TolQ-TolR proteins energize TolA and share homologies with the flagellar motor proteins MotA-MotB. Mol Microbiol 42, 795–807 (2001).

2. C. A. Santos, R. Janissen, M. A. Toledo, L. L. Beloti, A. R. Azzoni, M. A. Cotta, A. P. Souza, Characterization of the TolB-Pal trans-envelope complex from Xylella fastidiosa reveals a dynamic and coordinated protein expression profile during the biofilm development process. Biochim Biophys Acta 1854, 1372–1381 (2015).

3. R. Derouiche, H. Bénédetti, J. C. Lazzaroni, C. Lazdunski, R. Lloubès, Protein complex within Escherichia coli inner membrane. TolA N-terminal domain interacts with TolQ and TolR proteins. J Biol Chem 270, 11078–11084 (1995).

4. E. Bouveret, R. Derouiche, A. Rigal, R. Lloubès, C. Lazdunski, H. Bénédetti, Peptidoglycan-associated lipoprotein-TolB interaction. A possible key to explaining the formation of contact sites between the inner and outer membranes of Escherichia coli. J Biol Chem 270, 11071–11077 (1995).

5. J. Szczepaniak, P. Holmes, K. Rajasekar, R. Kaminska, F. Samsudin, P. G. Inns, P. Rassam, S. Khalid, S. M. Murray, C. Redfield, C. Kleanthous, The lipoprotein Pal stabilises the bacterial outer membrane during constriction by a mobilisation-and-capture mechanism. Nat Commun 11, 1305 (2020).

6. A. J. F. Egan, Bacterial outer membrane constriction. Mol Microbiol 107, 676–687 (2018).

7. J. Szczepaniak, C. Press, C. Kleanthous, The multifarious roles of Tol-Pal in Gram-negative bacteria. FEMS Microbiol Rev 44, 490–506 (2020).

8. A. A. Yakhnina, T. G. Bernhardt, The Tol-Pal system is required for peptidoglycan-cleaving enzymes to complete bacterial cell division. Proc Natl Acad Sci U S A 117, 6777–6783 (2020).

9. P. Samire, B. Serrano, D. Duché, E. Lemarié, R. Lloubès, L. Houot, Decoupling Filamentous Phage Uptake and Energy of the TolQRA Motor in Escherichia coli. J Bacteriol 202, (2020).

10. J. C. Lazzaroni, J. F. Dubuisson, A. Vianney, The Tol proteins of Escherichia coli and their involvement in the translocation of group A colicins. Biochimie 84, 391–397 (2002).

11. P. Germon, M. C. Ray, A. Vianney, J. C. Lazzaroni, Energy-dependent conformational change in the TolA protein of Escherichia coli involves its N-terminal domain, TolQ, and TolR. J Bacteriol 183, 4110–4114 (2001).

12. J. A. Wojdyla, E. Cutts, R. Kaminska, G. Papadakos, J. T. Hopper, P. J. Stansfeld, D. Staunton, C. V. Robinson, C. Kleanthous, Structure and function of the Escherichia coli Tol-Pal stator protein TolR. J Biol Chem 290, 26675–26687 (2015).

13. K. Kampfenkel, V. Braun, Membrane topologies of the TolQ and TolR proteins of Escherichia coli: inactivation of TolQ by a missense mutation in the proposed first transmembrane segment. J Bacteriol 175, 4485–4491 (1993).

14. A. Vianney, T. M. Lewin, W. F. Beyer, Jr., J. C. Lazzaroni, R. Portalier, R. E. Webster, Membrane topology and mutational analysis of the TolQ protein of Escherichia coli required for the uptake of macromolecules and cell envelope integrity. J Bacteriol 176, 822–829 (1994).

15. V. Braun, C. Herrmann, Point mutations in transmembrane helices 2 and 3 of ExbB and TolQ affect their activities in Escherichia coli K-12. J Bacteriol 186, 4402–4406 (2004).

16. J. C. Lazzaroni, P. Germon, M. C. Ray, A. Vianney, The Tol proteins of Escherichia coli and their involvement in the uptake of biomolecules and outer membrane stability. FEMS Microbiol Lett 177, 191–197 (1999).

17. X. Y. Zhang, E. L. Goemaere, R. Thomé, M. Gavioli, E. Cascales, R. Lloubès, Mapping the interactions between escherichia coli tol subunits: rotation of the TolR transmembrane helix. J Biol Chem 284, 4275–4282 (2009).

18. X. Y. Zhang, E. L. Goemaere, N. Seddiki, H. Célia, M. Gavioli, E. Cascales, R. Lloubes, Mapping the interactions between Escherichia coli TolQ transmembrane segments. J Biol Chem 286, 11756–11764 (2011).

19. J. C. Lazzaroni, A. Vianney, J. L. Popot, H. Bénédetti, F. Samatey, C. Lazdunski, R. Portalier, V. Géli, Transmembrane alpha-helix interactions are required for the functional assembly of the Escherichia coli Tol complex. J Mol Biol 246, 1–7 (1995).

20. P. Germon, T. Clavel, A. Vianney, R. Portalier, J. C. Lazzaroni, Mutational analysis of the Escherichia coli K-12 TolA N-terminal region and characterization of its TolQ-interacting domain by genetic suppression. J Bacteriol 180, 6433–6439 (1998).

21. L. Journet, A. Rigal, C. Lazdunski, H. Bénédetti, Role of TolR N-terminal, central, and C-terminal domains in dimerization and interaction with TolA and tolQ. J Bacteriol 181, 4476–4484 (1999).

22. R. Masilamani, M. B. Cian, Z. D. Dalebroux, Salmonella Tol-Pal Reduces Outer Membrane Glycerophospholipid Levels for Envelope Homeostasis and Survival during Bacteremia. Infect Immun 86, (2018).

23. E. L. Goemaere, E. Cascales, R. Lloubès, "Mutational analyses define helix organization and key residues of a bacterial membrane energy-transducing complex" in J Mol Biol (England, 2007), vol. 366, pp. 1424–1436.

24. J. F. Dubuisson, A. Vianney, N. Hugouvieux-Cotte-Pattat, J. C. Lazzaroni, Tol-Pal proteins are critical cell envelope components of Erwinia chrysanthemi affecting cell morphology and virulence. Microbiology (Reading) 151, 3337–3347 (2005).

25. J. A. Gaspar, J. A. Thomas, C. L. Marolda, M. A. Valvano, Surface expression of O-specific lipopolysaccharide in Escherichia coli requires the function of the TolA protein. Mol Microbiol 38, 262–275 (2000).

26. H. Hirakawa, K. Suzue, K. Kurabayashi, H. Tomita, The Tol-Pal System of Uropathogenic Escherichia coli Is Responsible for Optimal Internalization Into and Aggregation Within Bladder Epithelial Cells, Colonization of the Urinary Tract of Mice, and Bacterial Motility. Front Microbiol 10, 1827 (2019).

27. H. Abdelhamed, J. Lu, M. L. Lawrence, A. Karsi, Involvement of tolQ and tolR genes in Edwardsiella ictaluri virulence. Microb Pathog 100, 90–94 (2016).

28. Q. Li, Z. Li, X. Fei, Y. C. Tian, G. D. Zhou, Y. H. Hu, S. F. Wang, H. Y. Shi, The role of TolA, TolB, and TolR in cell morphology, OMVs production, and virulence of Salmonella Choleraesuis. Amb Express 12, (2022).

29. M. Witty, C. Sanz, A. Shah, J. G. Grossmann, K. Mizuguchi, R. N. Perham, B. Luisi, Structure of the periplasmic domain of Pseudomonas aeruginosa TolA: evidence for an evolutionary relationship with the TonB transporter protein. Embo j 21, 4207–4218 (2002).

30. C. Deprez, R. Lloubès, M. Gavioli, D. Marion, F. Guerlesquin, L. Blanchard, Solution structure of the E.coli TolA C-terminal domain reveals conformational changes upon binding to the phage g3p N-terminal domain. J Mol Biol 346, 1047–1057 (2005).

31. C. G. Ford, S. Kolappan, H. T. Phan, M. K. Waldor, H. C. Winther-Larsen, L. Craig, Crystal structures of a CTXphi pIII domain unbound and in complex with a Vibrio cholerae TolA domain reveal novel interaction interfaces. J Biol Chem 287, 36258–36272 (2012).

32. C. Li, Y. Zhang, M. Vankemmelbeke, O. Hecht, F. S. Aleanizy, C. Macdonald, G. R. Moore, R. James, C. N. Penfold, Structural evidence that colicin A protein binds to a novel binding site of TolA protein in Escherichia coli periplasm. J Biol Chem 287, 19048–19057 (2012).

33. L. M. Parsons, A. Grishaev, A. Bax, The periplasmic domain of TolR from Haemophilus influenzae forms a dimer with a large hydrophobic groove: NMR solution structure and comparison to SAXS data. Biochemistry 47, 3131–3142 (2008).

34. D. P. Williams-Jones, M. N. Webby, C. E. Press, J. M. Gradon, S. R. Armstrong, J. Szczepaniak, C. Kleanthous, Tunable force transduction through the Escherichia coli cell envelope. Proc Natl Acad Sci U S A 120, e2306707120 (2023).

35. S. Kojima, D. F. Blair, Conformational change in the stator of the bacterial flagellar motor. Biochemistry 40, 13041–13050 (2001).

36. H. Celia, N. Noinaj, S. D. Zakharov, E. Bordignon, I. Botos, M. Santamaria, T. J. Barnard, W. A. Cramer, R. Lloubes, S. K. Buchanan, Structural insight into the role of the Ton complex in energy transduction. Nature 538, 60–65 (2016).

37. H. Celia, I. Botos, X. Ni, T. Fox, N. De Val, R. Lloubes, J. Jiang, S. K. Buchanan, Cryo-EM structure of the bacterial Ton motor subcomplex ExbB-ExbD provides information on structure and stoichiometry. Commun Biol 2, 358 (2019).

38. M. Santiveri, A. Roa-Eguiara, C. Kühne, N. Wadhwa, H. Hu, H. C. Berg, M. Erhardt, N. M. I. Taylor, Structure and Function of Stator Units of the Bacterial Flagellar Motor. Cell 183, 244–257.e216 (2020).

39. J. C. Deme, S. Johnson, O. Vickery, A. Aron, H. Monkhouse, T. Griffiths, R. H. James, B. C. Berks, J. W. Coulton, P. J. Stansfeld, S. M. Lea, Structures of the stator complex that drives rotation of the bacterial flagellum. Nat Microbiol 5, 1553–1564 (2020).

40. Y. W. Lai, P. Ridone, G. Peralta, M. M. Tanaka, M. A. B. Baker, Evolution of the Stator Elements of Rotary Prokaryote Motors. J Bacteriol 202, (2020).

41. M. F. El-Badawy, F. I. Abou-Elazm, M. S. Omar, M. E. El-Naggar, I. A. Maghrabi, The First Saudi Study Investigating the Plasmid-borne Aminoglycoside and Sulfonamide Resistance among Acinetobacter baumannii Clinical Isolates Genotyped by RAPD-PCR: the Declaration of a Novel Allelic Variant Called aac(6’)-SL and Three Novel Mutations in the sul1 Gene in the Acinetobacter Plasmid (s). Infect Drug Resist 14, 4739–4756 (2021).

42. C. L. Williams, H. M. Neu, Y. A. Alamneh, R. M. Reddinger, A. C. Jacobs, S. Singh, R. Abu-Taleb, S. L. J. Michel, D. V. Zurawski, D. S. Merrell, Characterization of Acinetobacter baumannii Copper Resistance Reveals a Role in Virulence. Front Microbiol 11, 16 (2020).

43. F. Runci, V. Gentile, E. Frangipani, G. Rampioni, L. Leoni, M. Lucidi, D. Visaggio, G. Harris, W. Chen, J. Stahl, B. Averhoff, P. Visca, Contribution of Active Iron Uptake to Acinetobacter baumannii Pathogenicity. Infect Immun 87, (2019).

44. M. S. Mulani, E. E. Kamble, S. N. Kumkar, M. S. Tawre, K. R. Pardesi, Emerging Strategies to Combat ESKAPE Pathogens in the Era of Antimicrobial Resistance: A Review. Frontiers in Microbiology 10, 1–24 (2019).

45. M. van Kempen, S. S. Kim, C. Tumescheit, M. Mirdita, J. Lee, C. L. M. Gilchrist, J. Söding, M. Steinegger, Fast and accurate protein structure search with Foldseek. Nat Biotechnol, (2023).

46. T. Nishikino, N. Takekawa, D. P. Tran, J. I. Kishikawa, M. Hirose, S. Onoe, S. Kojima, M. Homma, A. Kitao, T. Kato, K. Imada, Structure of MotA, a flagellar stator protein, from hyperthermophile. Biochem Biophys Res Commun 631, 78–85 (2022).

47. E. L. Goemaere, A. Devert, R. Lloubès, E. Cascales, Movements of the TolR C-terminal domain depend on TolQR ionizable key residues and regulate activity of the Tol complex. J Biol Chem 282, 17749–17757 (2007).

48. S. Rao, G. Klesse, P. J. Stansfeld, S. J. Tucker, M. S. P. Sansom, A heuristic derived from analysis of the ion channel structural proteome permits the rapid identification of hydrophobic gates. Proc Natl Acad Sci U S A 116, 13989–13995 (2019).

49. A. Punjani, D. J. Fleet, 3D variability analysis: Resolving continuous flexibility and discrete heterogeneity from single particle cryo-EM. Journal of Structural Biology 213, (2021).

50. E. Cascales, M. Gavioli, J. N. Sturgis, R. Lloubès, Proton motive force drives the interaction of the inner membrane TolA and outer membrane pal proteins in Escherichia coli. Mol Microbiol 38, 904–915 (2000).

51. J. Jumper, R. Evans, A. Pritzel, T. Green, M. Figurnov, O. Ronneberger, K. Tunyasuvunakool, R. Bates, A. Zidek, A. Potapenko, A. Bridgland, C. Meyer, S. A. A. Kohl, A. J. Ballard, A. Cowie, B. Romera-Paredes, S. Nikolov, R. Jain, J. Adler, …, D. Hassabis, Highly accurate protein structure prediction with AlphaFold. Nature 596, 583-+ (2021).

52. M. N. Webby, D. P. Williams-Jones, C. Press, C. Kleanthous, Force-Generation by the Trans-Envelope Tol-Pal System. Frontiers in Microbiology 13, (2022).

53. P. J. Loll, K. C. Grasty, D. D. Shultis, N. J. Guzman, M. C. Wiener, Discovery and structural characterization of the D-box, a conserved TonB motif that couples an inner- membrane motor to outer-membrane transport. J Biol Chem, 105723 (2024).

54. M. Zinke, M. Lejeune, A. Mechaly, B. Bardiaux, I. G. Boneca, P. Delepelaire, N. Izadi- Pruneyre, Ton motor conformational switch and peptidoglycan role in bacterial nutrient uptake. Nat Commun 15, 331 (2024).

55. A. C. Ratliff, S. K. Buchanan, H. Celia, The Ton Motor. Front Microbiol 13, 852955 (2022).

56. P. Demchick, A. L. Koch, The permeability of the wall fabric of Escherichia coli and Bacillus subtilis. Journal of Bacteriology 178, 768–773 (1996).

57. M. Rieu, R. Krutyholowa, N. M. I. Taylor, R. M. Berry, A new class of biological ion- driven rotary molecular motors with 5:2 symmetry. Frontiers in Microbiology 13, (2022).

58. J. J. Kerrigan, Q. Xie, R. S. Ames, Q. Lu, Production of protein complexes via co- expression. Protein Expression and Purification 75, 1–14 (2011).

59. A. Olerinyova, A. Sonn-Segev, J. Gault, C. Eichmann, J. Schimpf, A. H. Kopf, L. S. P. Rudden, D. Ashkinadze, R. Bomba, L. Frey, J. Greenwald, M. T. Degiacomi, R. Steinhilper, J. A. Killian, T. Friedrich, R. Riek, W. B. Struwe, P. Kukura, Mass Photometry of Membrane Proteins. Chem 7, 224–236 (2021).

60. A. Punjani, J. L. Rubinstein, D. J. Fleet, M. A. Brubaker, cryoSPARC: algorithms for rapid unsupervised cryo-EM structure determination. Nat Methods 14, 290–296 (2017).

61. A. Punjani, H. Zhang, D. J. Fleet, Non-uniform refinement: adaptive regularization improves single-particle cryo-EM reconstruction. Nat Methods 17, 1214–1221 (2020).

62. E. F. Pettersen, T. D. Goddard, C. C. Huang, G. S. Couch, D. M. Greenblatt, E. C. Meng, T. E. Ferrin, UCSF Chimera--a visualization system for exploratory research and analysis. J Comput Chem 25, 1605–1612 (2004).

63. P. Emsley, K. Cowtan, Coot: model-building tools for molecular graphics. Acta Crystallogr D Biol Crystallogr 60, 2126–2132 (2004).

64. E. Krissinel, K. Henrick, Secondary-structure matching (SSM), a new tool for fast protein structure alignment in three dimensions. Acta Crystallogr D Biol Crystallogr 60, 2256–2268 (2004).

65. D. Liebschner, P. V. Afonine, M. L. Baker, G. Bunkóczi, V. B. Chen, T. I. Croll, B. Hintze, L. W. Hung, S. Jain, A. J. McCoy, N. W. Moriarty, R. D. Oeffner, B. K. Poon, M. G. Prisant, R. J. Read, J. S. Richardson, D. C. Richardson, M. D. Sammito, O. V. Sobolev, …, P. D. Adams, Macromolecular structure determination using X-rays, neutrons and electrons: recent developments in Phenix. Acta Crystallogr D Struct Biol 75, 861–877 (2019).

66. K. Yamashita, C. M. Palmer, T. Burnley, G. N. Murshudov, Cryo-EM single-particle structure refinement and map calculation using Servalcat. Acta Crystallogr D Struct Biol 77, 1282–1291 (2021).

67. A. Brown, F. Long, R. A. Nicholls, J. Toots, P. Emsley, G. Murshudov, Tools for macromolecular model building and refinement into electron cryo-microscopy reconstructions. Acta Crystallogr D Biol Crystallogr 71, 136–153 (2015).

68. P. V. Afonine, B. K. Poon, R. J. Read, O. V. Sobolev, T. C. Terwilliger, A. Urzhumtsev, P. D. Adams, Real-space refinement in PHENIX for cryo-EM and crystallography. Acta Crystallogr D Struct Biol 74, 531–544 (2018).

69. L. L. C. Schrödinger, "The PyMOL Molecular Graphics System, Version 2.3." (2020; http://www.pymol.org/pymol).

70. P. C. Hsu, F. Samsudin, J. Shearer, S. Khalid, It Is Complicated: Curvature, Diffusion, and Lipid Sorting within the Two Membranes of Escherichia coli. J Phys Chem Lett 8, 5513–5518 (2017).

71. E. L. Wu, P. J. Fleming, M. S. Yeom, G. Widmalm, J. B. Klauda, K. G. Fleming, W. Im, E. coli outer membrane and interactions with OmpLA. Biophys J 106, 2493–2502 (2014).

72. V. W. Rowlett, V. Mallampalli, A. Karlstaedt, W. Dowhan, H. Taegtmeyer, W. Margolin, H. Vitrac, Impact of Membrane Phospholipid Alterations in Escherichia coli on Cellular Function and Bacterial Stress Adaptation. J Bacteriol 199, (2017).

73. S. Jo, J. B. Lim, J. B. Klauda, W. Im, CHARMM-GUI Membrane Builder for mixed bilayers and its application to yeast membranes. Biophys J 97, 50–58 (2009).

74. E. L. Wu, X. Cheng, S. Jo, H. Rui, K. C. Song, E. M. Dávila-Contreras, Y. Qi, J. Lee, V. Monje-Galvan, R. M. Venable, J. B. Klauda, W. Im, CHARMM-GUI Membrane Builder toward realistic biological membrane simulations. J Comput Chem 35, 1997–2004 (2014).

75. S. Jo, T. Kim, W. Im, Automated builder and database of protein/membrane complexes for molecular dynamics simulations. PLoS One 2, e880 (2007).

76. J. C. Phillips, D. J. Hardy, J. D. C. Maia, J. E. Stone, J. V. Ribeiro, R. C. Bernardi, R. Buch, G. Fiorin, J. Hénin, W. Jiang, R. McGreevy, M. C. R. Melo, B. K. Radak, R. D. Skeel, A. Singharoy, Y. Wang, B. Roux, A. Aksimentiev, Z. Luthey-Schulten, …, E. Tajkhorshid, Scalable molecular dynamics on CPU and GPU architectures with NAMD. J Chem Phys 153, 044130 (2020).

77. J. Huang, S. Rauscher, G. Nawrocki, T. Ran, M. Feig, B. L. de Groot, H. Grubmüller, A. D. MacKerell, Jr., CHARMM36m: an improved force field for folded and intrinsically disordered proteins. Nat Methods 14, 71–73 (2017).

78. J. B. Klauda, R. M. Venable, J. A. Freites, J. W. O’Connor, D. J. Tobias, C. Mondragon- Ramirez, I. Vorobyov, A. D. MacKerell, Jr., R. W. Pastor, Update of the CHARMM all- atom additive force field for lipids: validation on six lipid types. J Phys Chem B 114, 7830–7843 (2010).

79. U. Essmann, L. Perera, M. L. Berkowitz, T. Darden, H. Lee, L. G. Pedersen, A SMOOTH PARTICLE MESH EWALD METHOD. Journal of Chemical Physics 103, 8577–8593 (1995).

80. W. Humphrey, A. Dalke, K. Schulten, VMD: visual molecular dynamics. J Mol Graph 14, 33–38, 27-38 (1996).

81. N. Michaud-Agrawal, E. J. Denning, T. B. Woolf, O. Beckstein, MDAnalysis: a toolkit for the analysis of molecular dynamics simulations. J Comput Chem 32, 2319–2327 (2011).

82. O. S. Smart, J. G. Neduvelil, X. Wang, B. A. Wallace, M. S. Sansom, HOLE: a program for the analysis of the pore dimensions of ion channel structural models. J Mol Graph 14, 354–360, 376 (1996).

83. M. Mirdita, K. Schütze, Y. Moriwaki, L. Heo, S. Ovchinnikov, M. Steinegger, ColabFold: making protein folding accessible to all. Nat Methods 19, 679–682 (2022).

84. E. C. Meng, T. D. Goddard, E. F. Pettersen, G. S. Couch, Z. J. Pearson, J. H. Morris, T. E. Ferrin, UCSF ChimeraX: Tools for structure building and analysis. Protein Sci 32, e4792 (2023).

85. J. D. Hunter, Matplotlib: A 2D graphics environment. Computing in Science & Engineering 9, 90–95 (2007).

86. R. A. Laskowski, J. Jabłońska, L. Pravda, R. S. Vařeková, J. M. Thornton, PDBsum: Structural summaries of PDB entries. Protein Sci 27, 129–134 (2018).

87. X. Robert, P. Gouet, Deciphering key features in protein structures with the new ENDscript server. Nucleic Acids Research 42, W320–W324 (2014).

