## Supplemental material for "Structural architecture of TolQ-TolR inner membrane protein complex from opportunistic pathogen *Acinetobacter baumannii*"

**This PDF file includes:**

Figs. S1 to S17
Tables S1 to S3

**Other Supplementary Materials for this manuscript include the following:**

Movies S1 to S5

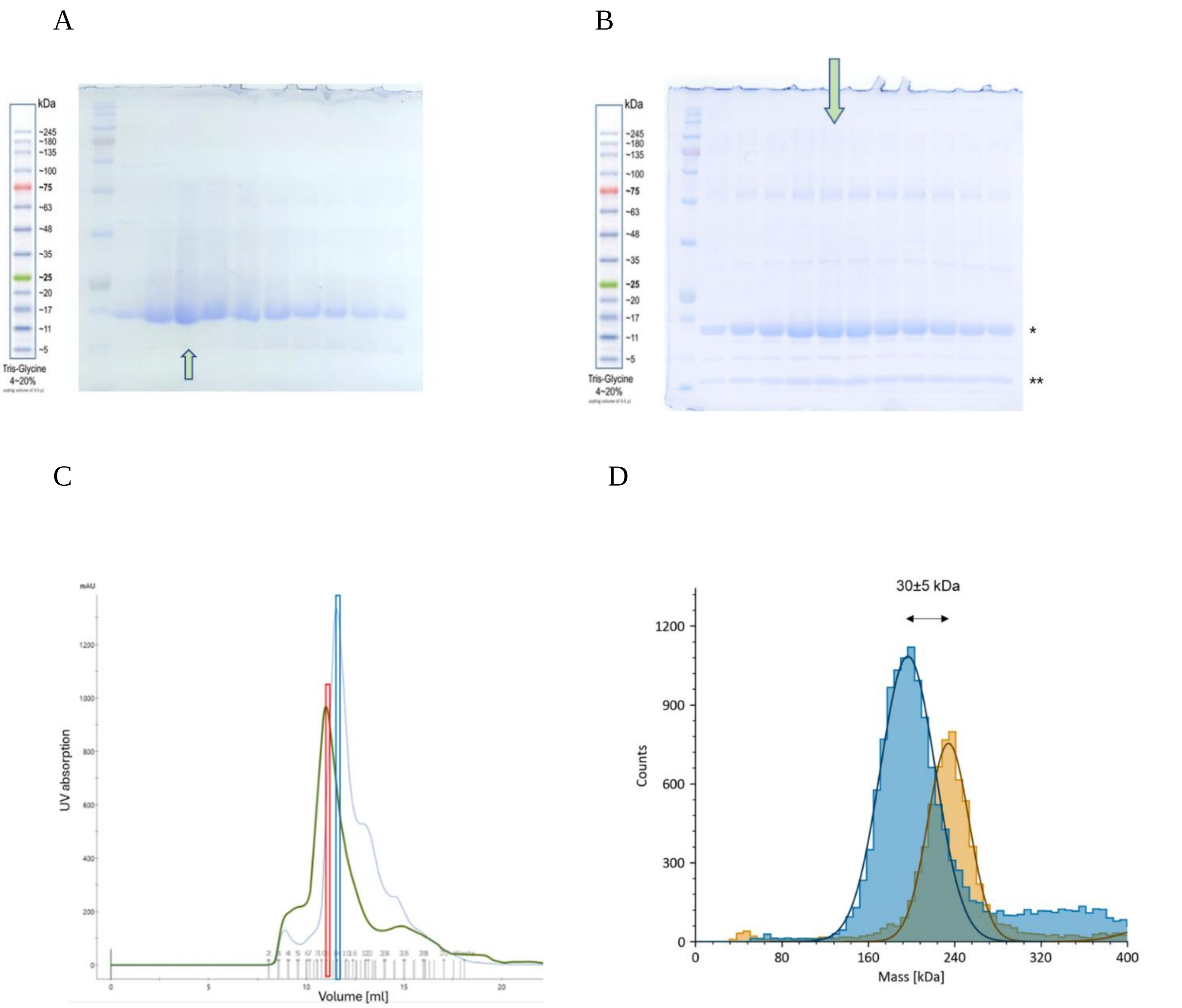


**Fig. S1.**

The quality assays of the apo *Ac*TolQ and *Ac*TolQ-TolR samples taken for cryo-EM analysis. (A) The SDS PAGE gel of purified *Ac*TolQ during size exclusion (SEC). The green arrow indicates the fraction taken further for analysis. (B) The SDS PAGE gel of purified *Ac*TolQ-TolR during size exclusion. A star indicates the *Ac*TolQ protein band. A double star indicates the *Ac*TolR band. The green arrow indicates the fraction taken further for analysis. (C) SEC profile of *Ac*TolQ (blue) and *Ac*TolQ-TolR (green). Red and bright blues bars indicate fractions taken for analysis from *Ac*TolQ and *Ac*TolQ-TolR samples, respectively (D) Molecular mass distribution histograms of *Ac*TolQ (blue) and *Ac*TolQ-TolR (orange) samples using mass photometry.

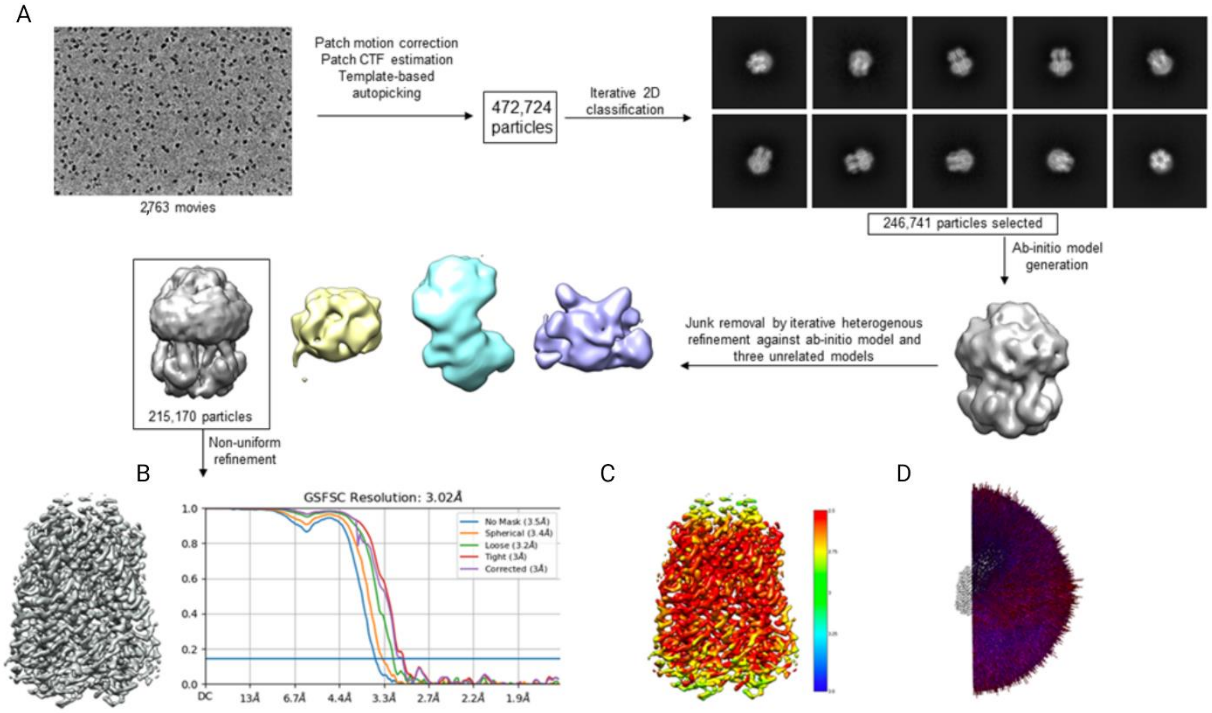


**Fig. S2.**

*Ac*TolQ single particle reconstruction flow chart. (A) Data collection and 2D classification. (B) Best resolving volume and refinement, and 3D Fourier shell correlation (FSC) curves for *Ac*TolQ. (C) Cryo-EM density map of *Ac*TolQ colored by local resolution (in Å). (D) Euler angular distribution plotting.

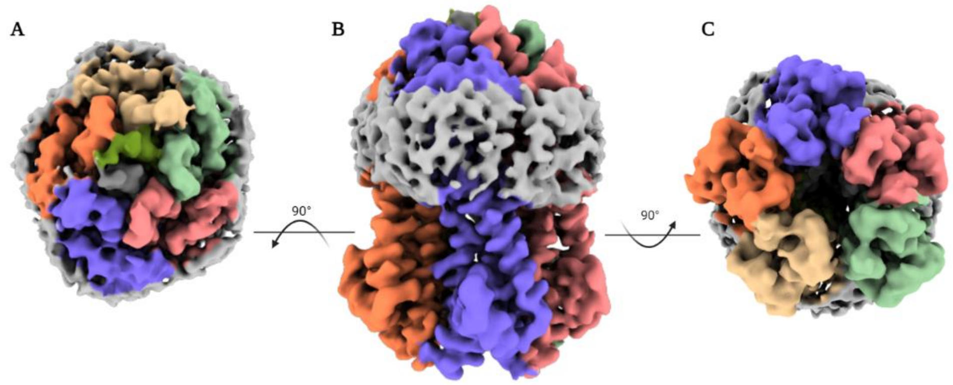


**Fig. S3.**

Low contour electron density map with amphipol TMH stabilizing region. (A) Periplasmic view. (B) Side view. (C) Cytoplasmic view.

Interface #5 in 06262.pdb//B:A

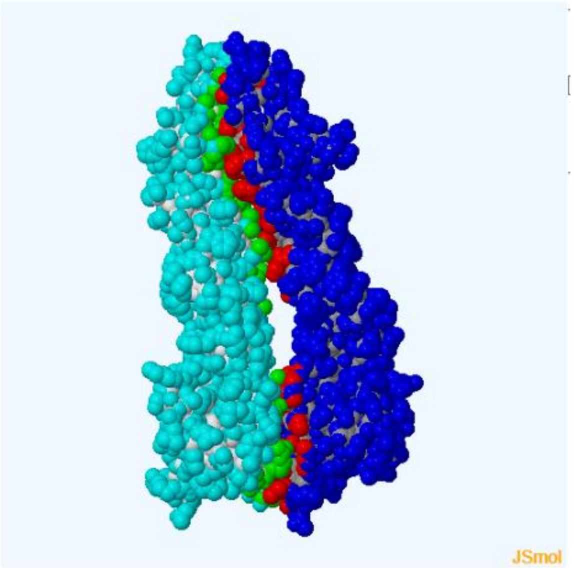


**Fig. S4.**

Oligomerization interface (green/red) between chain 1 (light blue) and chain 2 (dark blue) *Ac*TolQ protomers calculated using PISA.

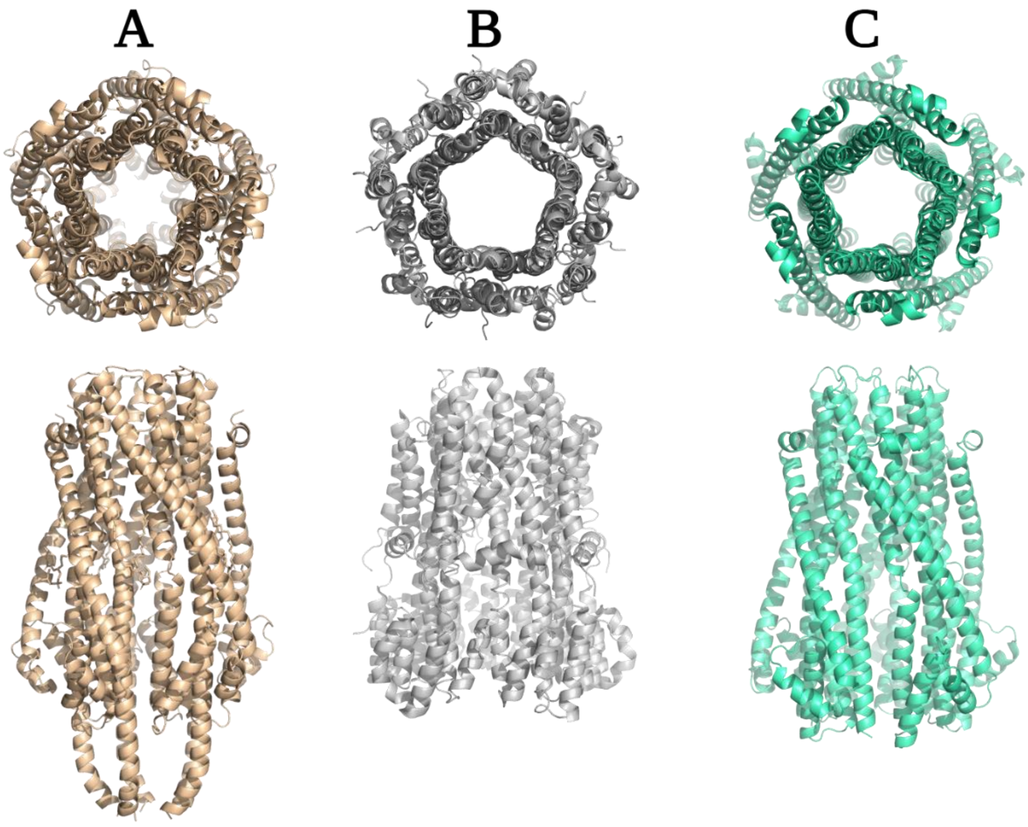


**Fig. S5.**

Symmetrical pore formation by five protomers of (A) ExbB (PDB:6YE4), (B) MotA (PDB:8GQY), and (C) TolQ (PDB:9AVI) proteins viewed from the periplasmic side (top row) and a side view (bottom row).

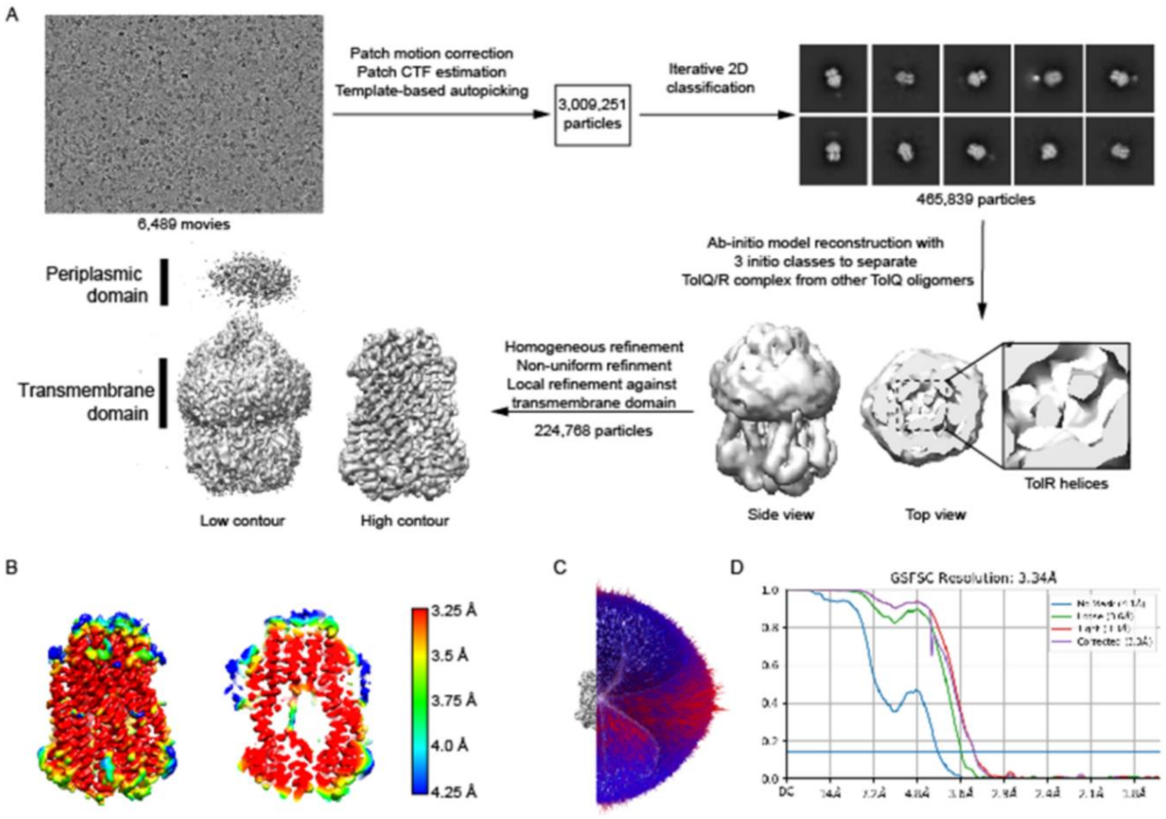


**Fig. S6.**

*Ac*TolQ-TolR single particle reconstruction flow chart. (A) Data collection, 2D classification, *ab initio* model building and refinement. (B) Cryo-EM density map of *Ac*TolQ-TolR colored by local resolution (in Å). (C) Euler angular distribution plotting. (D) 3D Fourier shell correlation (FSC) curves for *Ac*TolQ-TolR.

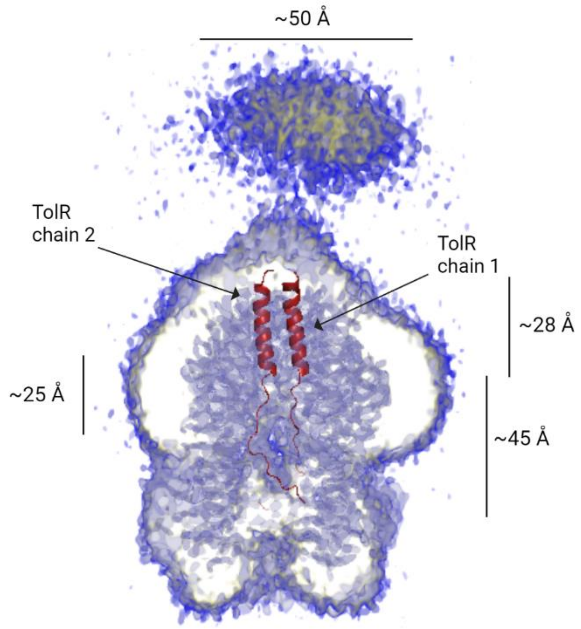


**Fig. S7.**

Low contour electron density map with periplasmic domain of TolR seen at periplasmic side of the TolQ-TolR complex.

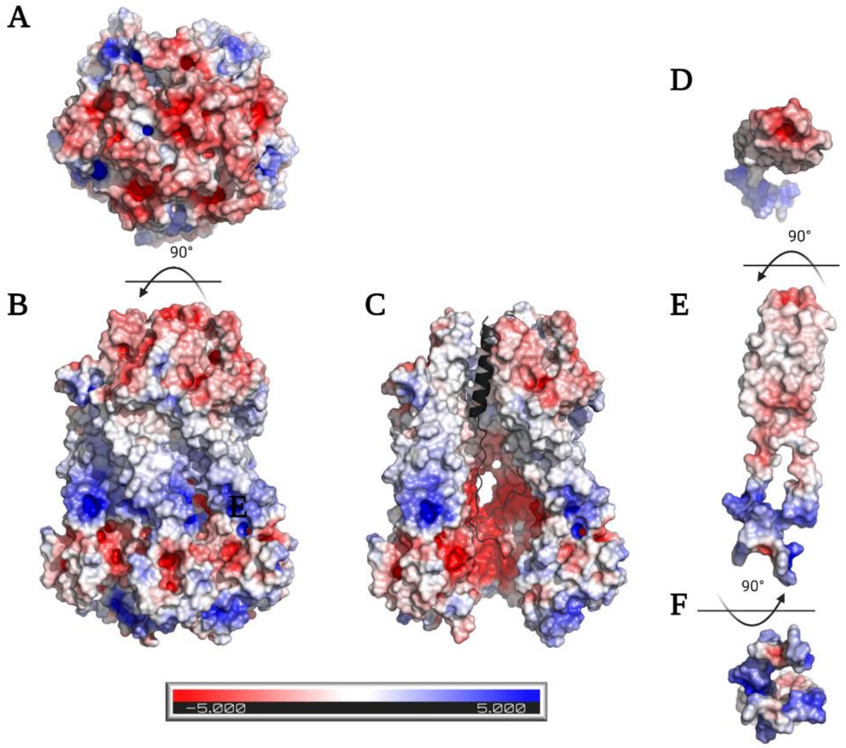


**Fig. S8.**

Electrostatic surface representation. The surface electrostatic potential of the *Ac*TolQ-TolR complex was calculated by APBS (Pymol). The potential is given with the negative (red) and positive (blue) contour levels ranging from −5.0 to +5.0 kBT respectively. (A) *Ac*TolQ-TolR complex periplasmic view. (B) *Ac*TolQ-TolR complex side view. (C) Electrostatic surface representation of the cytoplasmic chamber and transmembrane channel inner view formed by *Ac*TolQ protomers. One *Ac*TolQ protomer was eliminated for the proper view. (D) The *Ac*TolR dimer's electrostatic surface representation from the periplasmic top view (E) The *Ac*TolR dimer's electrostatic surface representation from the side view (F) The *Ac*TolR dimer's electrostatic surface representation from the cytoplasmic bottom view.

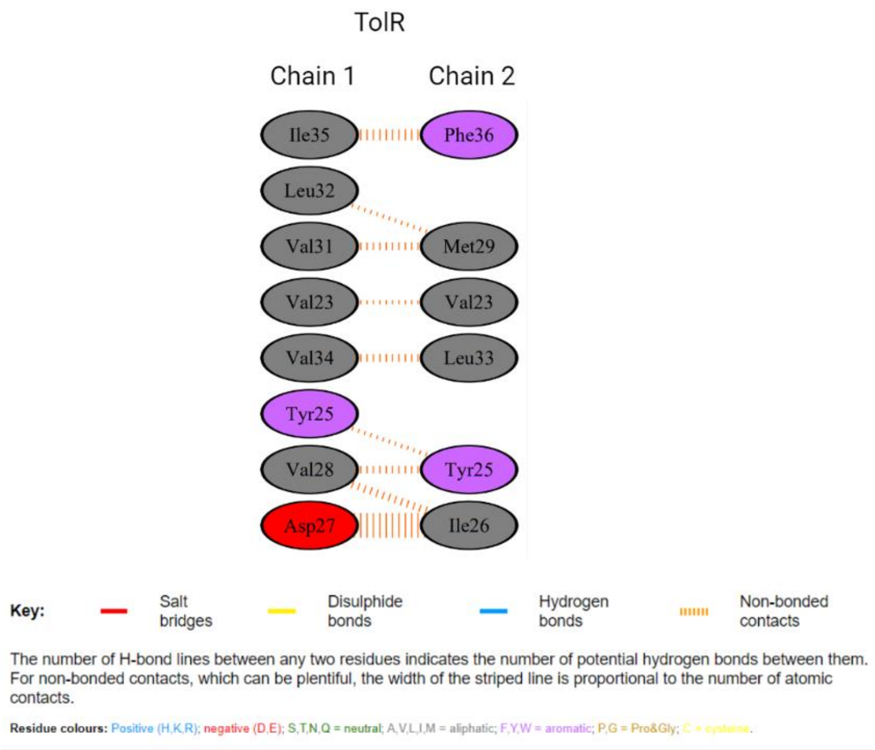


**Fig. S9.**

Interactions at the *Ac*TolR dimer interface. Residues are colored by type. This figure was generated using PDBsum web server.

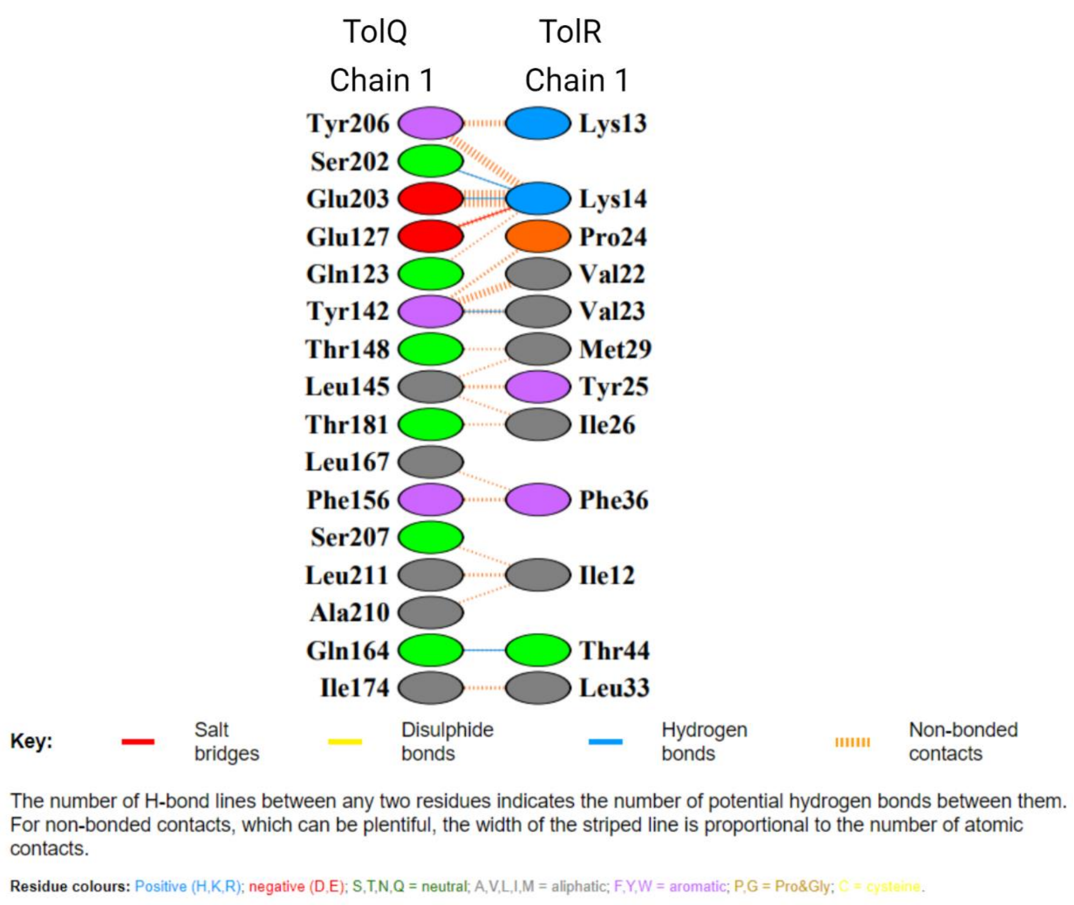


**Fig. S10.**

Interactions at the *Ac*TolQ chain 1 with *Ac*TolR chain 1 interface. Residues are colored by type. This figure was generated using PDBsum web server.

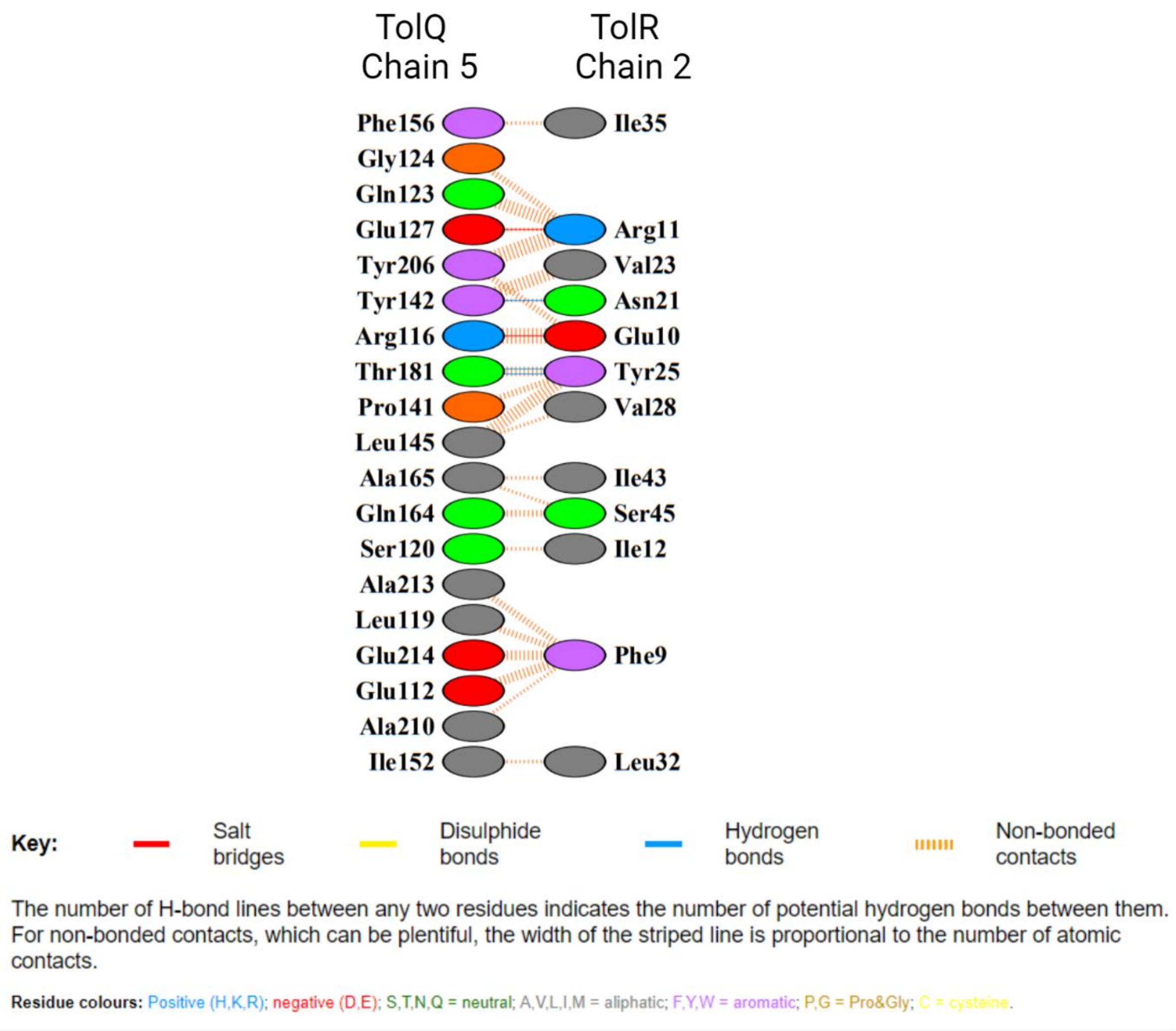


**Fig. S11.**

Interactions at the *Ac*TolQ chain 5 and *Ac*TolR chain 2 interfaces. Residues were colored by type. This figure was generated using PDBsum web server.

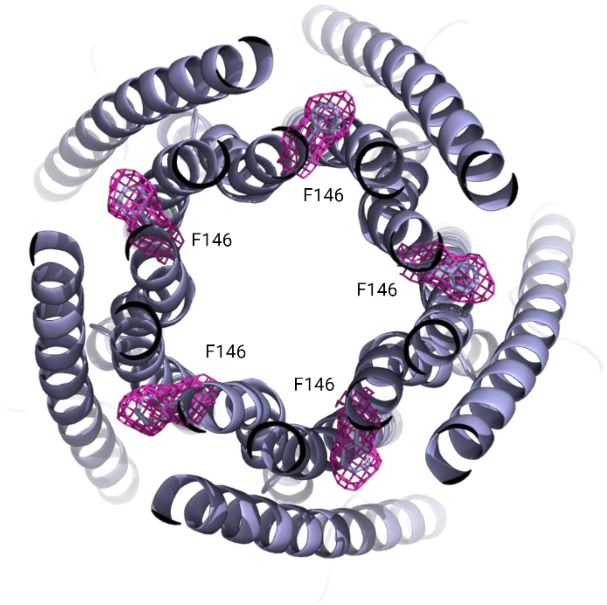


**Fig. S12.**

Slab view (from the periplasm) of the side chain orientation of $F^{146}$ in apo *Ac*TolQ with electron density depicted in magenta.

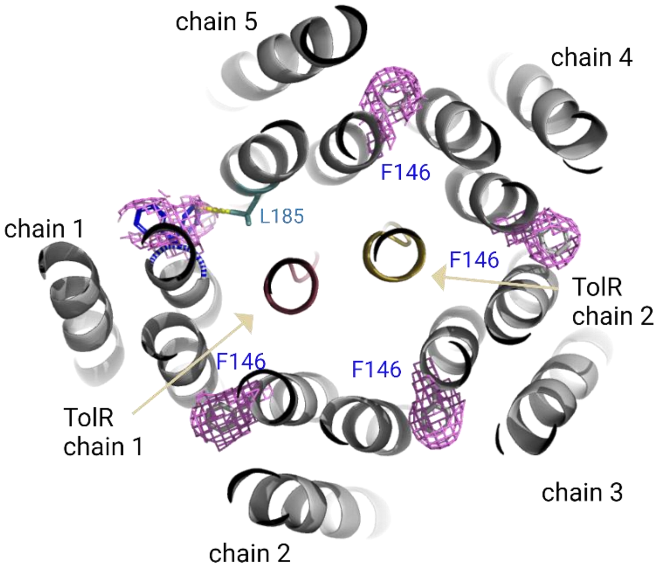


**Fig. S13.**

Slab view (from the periplasm) of the side chain orientation of $F^{146}$ in *Ac*TolQ when in complex with *Ac*TolR with electron density depicted in magenta.

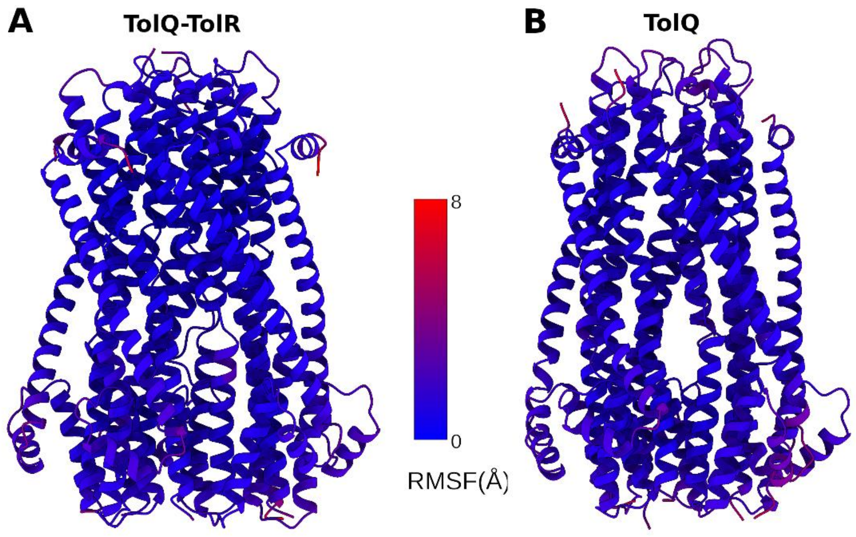


**Fig. S14.**

RMSF from MD simulations. (A) *Ac*TolQ-TolR and (B) *Ac*TolQ structures.

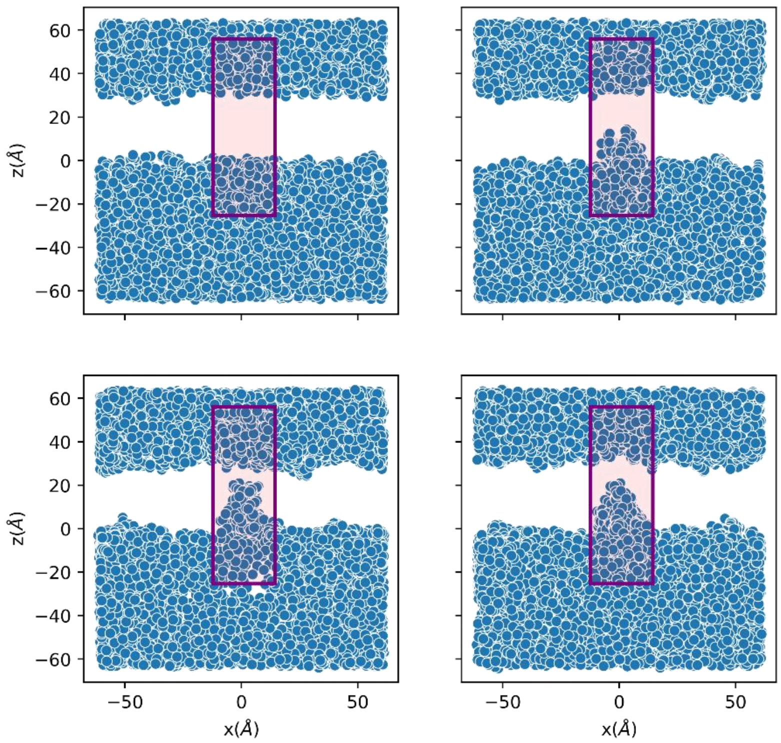


**Fig. S15.**

Ion dynamics of $K^+$ ions from MD simulations of TolQ-TolR structures. Selected ions are highlighted using their xz projections from the trajectories. The boxed shaded area (magenta) represents the TolR dimer positions and region of putative ion/proton flow. The upper left panel highlights an ion (control) that never enters the pore region of the TolQ-TolR. The other panel indicates three other $K^+$ ions that enter the pore region from the cytoplasmic side, however, never form a continuous path highlighting selective rejection from the TolR dimer upper part.

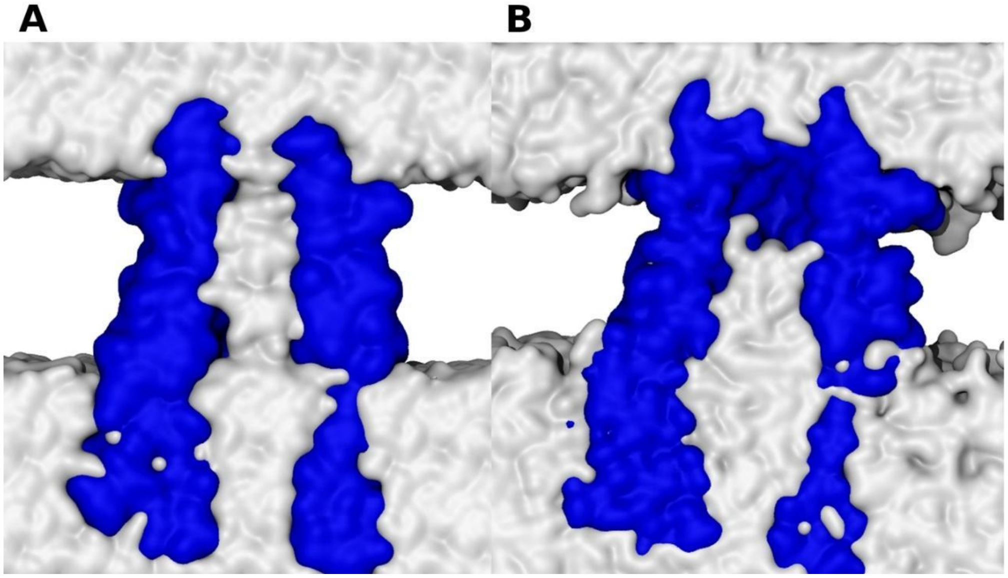

**Fig. S16.**

Dewetting of the TolQ structure. (A) Beginning of the simulation with generated pore water, (B) dewetted structure after simulation. The dewetting occurs rapidly and the pore remains dewetted throughout the micro-second-long simulation. Blue surface represents cross-sectional view of the $TolQ_5$ complex and gray surface represents water.

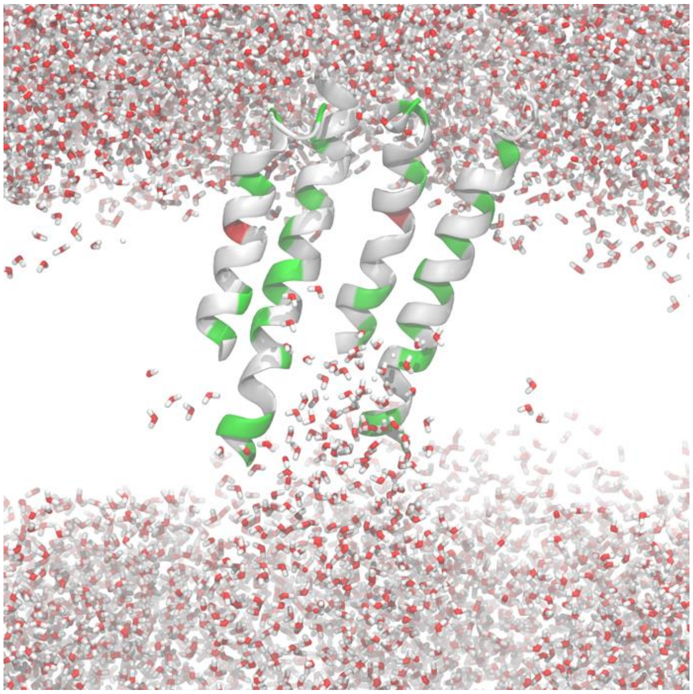

**Fig. S17.**

Residues forming the dewetting region in $TolQ_5$. Helices are shown in cartoon and water molecules in licorice. Helices from only two $TolQ_5$ protomers are shown for clarity. In the cartoon helices representation, nonpolar residues are colored in white, acidic residues are red, and polar residues are green.

**Table S1.**
Comparison of interface areas of TolQ protomers in apo form and when in complex with TolR using PISA

| Interfaces between chains | apo TolQ Interface area, Å$^2$ calculated by PISA | TolQ in complex with TolR Interface area, Å$^2$ calculated by PISA |
|---|---|---|
| Chains 1-2 (A and B) | 1228.6 | 1439.2 |
| Chains 2-3 (B and C) | 1233.7 | 1433.1 |
| Chains 3-4 (C and D) | 1231.6 | **754.8** |
| Chains 4-5 (D and E) | 1231.2 | 1435.6 |
| Chains 5-1 (E and A) | 1233.6 | **822.5** |
| Average | 1231.8 | 1177.0 |

**Table S2.**
Apo *Ac*TolQ cryo-EM data collection, refinement, and validation statistics

| | Data collection |
|---|---|
| Instrument | Titan Krios G2 |
| Detector | K3 Summit |
| Energy filter | Yes |
| Objective aperture | No |
| Nominal magnification | 105,000× |
| Data collection mode | Beam-Image Shift; 3×3 holes per movie |
| Frames per movie | 125 |
| Electron dose ($e^-/A^2$/frame) | ~0.5 |
| Exposure time (s/frame) | 0.035 |
| Super-resolution mode | Yes |
| Detector pixel size (Å) | 0.833 |
| Data pixel size (Å) | 0.417 |
| Movies acquired | 2,718 |
| | Reconstruction |
| Molecular weight (kDa) | 25.37 × 5 |
| Reconstruction symmetry | C5 |
| Particles used in refinement | 215,170 |
| Resolution $FSC_{0.143}$ (Å) | 3.02 |
| | Refinement and validation |
| Non-hydrogen atoms | 8,535 |
| Protein residues | 1,080 |
| Ligands | 0 |
| RMSD bond lengths (Å) | 0.007 |
| RMSD bond angles (°) | 1.447 |
| Model-to-map FSC (all) | 0.7571 |
| MolProbity score | 2.57 |

| | |
|---|---|
| Clashscore (all atom) | 9.05 |
| Poor rotamers (%) | 6.25 |
| Ramachandran (%) | |
| favored | 92.06 |
| allowed | 7.94 |
| outliers | 0 |

**Table S3.**

*Ac*TolQ-TolR cryo-EM data collection, refinement and validation statistics

| | Data collection |
|---|---|
| Instrument | Titan Krios G2 |
| Detector | K3 Summit |
| Energy filter | Yes |
| Objective aperture | No |
| Nominal magnification | 105,000× |
| Data collection mode | Beam-Image Shift; 3×3 holes per movie |
| Frames per movie | 100 |
| Electron dose ($e^-/A^2$/frame) | ~0.5 |
| Exposure time (s/frame) | 0.05 |
| Super-resolution mode | Yes |
| Detector pixel size (Å) | 0.834 |
| Data pixel size (Å) | 0.417 |
| Movies acquired | 6,489 |
| | Reconstruction |
| Molecular weight (kDa) | 25.37 × 5 + 16.73 × 2 |
| Reconstruction symmetry | C1 |
| Particles used in refinement | 224,768 |
| Resolution $FSC_{0.143}$ (Å) | 3.34 |
| | Refinement and validation |
| Non-hydrogen atoms | 9,091 |
| Protein residues | 1,173 |
| Ligands | 0 |
| RMSD bond lengths (Å) | 0.0141 |
| RMSD bond angles (°) | 1.6681 |
| Model-to-map FSC (all) | 0.707 |
| MolProbity score | 1.54 |
| Clashscore (all atom) | 8.31 |

| | |
|---|---|
| Poor rotamers (%) | 0.1 |
| Ramachandran (%) | |
| favored | 97.58 |
| allowed | 2.16 |
| outliers | 0.26 |

**Table S4.**
Simulation setup details of the systems.

| **System details** | **TolQ-TolR complex in membrane** | **TolQ complex in membrane** |
|---|---|---|
| Protein subunits | *Ac*TolQ-TolR (5-2) | *Ac*TolQ (5) |
| No of lipids | 400 | 400 |
| Composition | PPPE (312), PVPG (48), PVCL2 (24), POPA (16) | PPPE (312), PVPG (48), PVCL2 (24), POPA (16) |
| Dimension (Å) | ~ 127x127x129 | ~ 127x127x129 |
| Number of atoms | 189224 | 190982 |
| Number of waters | 39879 | 40641 |
| Pore water | - | 248 |
| No of $K^+$ | 222 | 228 |
| No of $Cl^-$ | 108 | 111 |

**Movie S1.**
3D variability assay, mode 0: cytoplasmic, side and slide views of AcTolQ-TolR densities.

**Movie S2.**
3D variability assay, mode 1: cytoplasmic, side and slide views of AcTolQ-TolR densities.

**Movie S3.**
3D variability assay, mode 2: cytoplasmic, side and slide views of AcTolQ-TolR densities.

**Movie S4.**
3D variability assay, mode 3: cytoplasmic, side and slide views of AcTolQ-TolR densities.

**Movie S5.**
3D variability assay, mode 4: cytoplasmic, side and slide views of AcTolQ-TolR densities.
